# Bacterial chemoreceptor signaling complexes control kinase activity by stabilizing the catalytic domain of CheA

**DOI:** 10.1101/2022.10.28.514197

**Authors:** Thomas Tran, Aruni P. K. K. Karunanayake Mudiyanselage, Stephen J. Eyles, Lynmarie K. Thompson

**Author notes:** Corresponding Author: Lynmarie K. Thompson, **Email:**. Authors contributed equally (Thomas Tran, Aruni P. K. K. Karunanayake Mudiyanselage). **Author Contributions:** T.T., A.K.M., S.J.E., and L.K.T. designed research; T.T. and A.K.M. performed research; T.T., A.K.M., S.J.E, and L.K.T. analyzed the data; T.T., A.K.M, and L.K.T. wrote the paper. **Competing Interest Statement:** The authors declare no competing interests.

## Abstract

Motile bacteria have a chemotaxis system that enables them to sense their environment and direct their swimming towards favorable conditions. Chemotaxis involves a signaling process in which ligand binding to the extracellular domain of the chemoreceptor alters the activity of the histidine kinase, CheA, bound ∼300 Å away to the distal cytoplasmic tip of the receptor, to initiate a phosphorylation cascade that controls flagellar rotation. The cytoplasmic domain of the receptor is thought to propagate this signal via changes in dynamics and/or stability, but it is unclear how these changes modulate the kinase activity of CheA. To address this question, we have used hydrogen deuterium exchange mass spectrometry to probe the structure and dynamics of CheA within functional signaling complexes of the *E. coli* aspartate receptor cytoplasmic fragment, CheA, and CheW. Our results reveal that stabilization of the P4 catalytic domain of CheA correlates with kinase activation. Furthermore, differences in activation of the kinase that occur during sensory adaptation depend on receptor destabilization of the P3 dimerization domain of CheA. Finally, hydrogen exchange properties of the P1 domain that bears the phosphorylated histidine identify the dimer interface of P1/P1’ in the CheA dimer and support an ordered sequential binding mechanism of catalysis, in which dimeric P1/P1’ has productive interactions with P4 only upon nucleotide binding. Thus stabilization/destabilization of domains is a key element of the mechanism of modulating CheA kinase activity in chemotaxis, and may play a role in the control of other kinases.

## Introduction

Signal transduction plays critical roles in cell viability by regulating pathways involved in cell growth, metabolism, and migration. Chemotaxis is a key signaling system that enables *E. coli* and other motile bacteria to migrate away from harmful conditions and towards favorable ones. Understanding the molecular mechanisms of bacterial chemotaxis proteins may facilitate disruption of their role in virulence processes such as biofilm formation (1). A central player in bacterial chemotaxis is the histidine kinase CheA, which is a cytoplasmic component of the large, membrane-bound protein arrays that sense the environment of the cell. In these arrays, multiple types of transmembrane chemoreceptors bind different ligands and this information is integrated to control the autophosphorylation activity of CheA. CheA catalyzes phosphoryl transfer to response regulators that ultimately control the direction of flagellar rotation. This sensory input enables cells to suppress tumbling in the presence of attractants, thus biasing their random walk towards favorable environments.

Although there are good structural models (2–4) of the *E. coli* sensor array, it is still unclear how the receptors control the activity of CheA. The transmembrane receptors form trimers-of-dimers (TOD) that are further associated via interactions at their membrane-distal cytoplasmic tips with the kinase CheA and a coupling protein CheW (Fig 1A). The core signaling unit (CSU, Fig 1A-B) consists of two receptor TODs bridged by a dimer of CheA and associated with two CheW monomers. These CSUs further associate into a supramolecular hexagonal array connected by rings of alternating CheA/CheW (Fig 1C) and rings of CheW. Sensing begins with the binding of ligands to the periplasmic domains of the chemoreceptors, for example binding of the attractant aspartate (Asp) to the Asp receptor. This binding causes a 2 Å piston displacement of a periplasmic domain helix connected to transmembrane helix 2 (5). Propagation of this signal through the cytoplasmic domain is thought to involve changes in dynamics (6, 7) within this partially disordered domain (8), that ultimately inhibit the kinase activity of CheA and phosphotransfer to the response regulator CheY that controls the direction of flagellar rotation.

**Figure 1.**
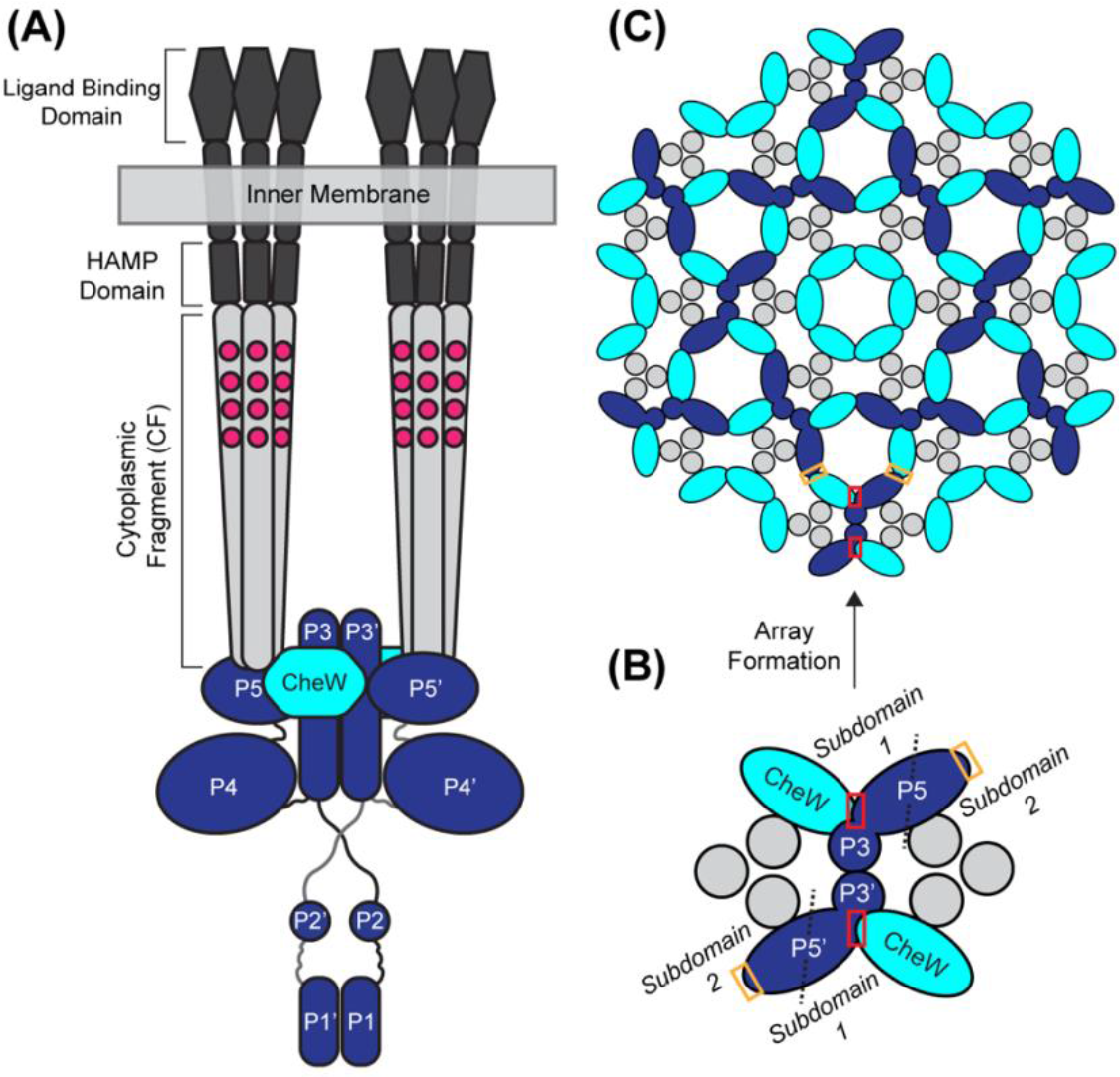
CheA Organization in Core Signaling Units and Signaling Arrays. (*A*) Core signaling unit (CSU) comprised of CheA dimer (blue, with its 5 domains labeled), two CheW (cyan), and two receptor trimer-of-dimers (CF in gray). Black lines represent linkers connecting the domains of CheA. (*B*) Bottom-up view of the CSU highlighting its interprotein interactions, including interface 1 (red box) within the CSU and interface 2 (orange box) that link the CSU into an array (51, 52). (*C*) Signaling array consisting of multiple CSUs connected by rings of alternating CheA and CheW, surrounding CheW-only rings. P1, P2 and P4 domains are not shown in B and C for simplicity.

Additional proteins mediate adaptation to an ongoing stimulus via methylation and demethylation of four glutamate residues within the cytoplasmic domain of the chemoreceptor, catalyzed by the CheR methyltransferase, and CheB methylesterase (also a response regulator phosphorylated by CheA). Receptor methylation shifts the array to the kinase-on state; receptor demethylation and ligand binding shift the array to the kinase-off state. However, it is not known how inhibition of CheA is achieved by ligand binding to the receptor or how this inhibition is relieved by methylation of the receptor.

The homodimeric CheA kinase (reviewed in (9)) is comprised of five domains (Fig 1A): P1 contains the phosphoryl-accepting histidine (H48 in *E. coli* CheA), P2 binds to response regulators (CheY and CheB), P3 mediates dimerization, P4 catalyzes the transfer of the γ-phosphate of ATP to His48, and P5 is the regulatory domain that interacts with CheW and the receptor to form the CSU, and interacts with CheW to form the CheA/CheW rings that connect the supramolecular signaling arrays. Phosphorylation occurs in *trans*, requiring interactions between P4 of one subunit and P1’ of the other subunit in the dimer. *E. coli* CheA has low kinase activity when not assembled into the signaling array. Kinase activity is enhanced ∼1000-fold-when CheA is bound to the kinase-on array (10, 11).

Understanding signaling mechanisms in this multiprotein complex requires assembly of functional complexes. For previous studies of the receptor structure and mechanism, our lab has assembled native-like complexes of the *E. coli* Asp receptor cytoplasmic fragment (CF) with CheA and CheW (12–14). Binding of the His-tagged CF to vesicles containing a Ni^2+^-chelating lipid headgroup in the presence of CheA and CheW leads to spontaneous assembly of complexes with both kinase and methylation activity (15) and the native hexagonal array structure (16). Hydrogen Deuterium Exchange Mass Spectrometry (HDX-MS) of CF in these vesicle-mediated functional complexes demonstrated that CF is partially disordered and that kinase activation is associated with reduced uptake, suggesting stabilization of CF. In particular, there was slower deuterium uptake in the protein interaction region of CF in its methylated, kinase-on state (represented by CF4Q, since glutamines mimic the methylated glutamates (17)) than in its normally kinase-off, unmethylated state (CF4E). We proposed that CheA is more tightly bound to CF4Q in kinase-on complexes, and that weaker binding interactions with CF4E allow it to revert to its kinase-off state (8).

Here we report an HDX-MS study of *E. coli* CheA in states with different extents of kinase activation, with the goal of determining what changes in structure and dynamics are involved in activating CheA. We measured HDX-MS of free CheA and CheA in functional signaling complexes. The vesicle-mediated functional complexes were assembled under conditions with 80-90% of CheA bound, in order to measure the properties of CheA within the complexes. We compared the hydrogen exchange of CheA in states associated with low kinase activity (CheA in solution and CheA in complexes with CF4E) to its hydrogen exchange in states associated with high kinase activity (CheA in complexes with CF4Q, with or without addition of a non-hydrolyzable ATP analog, β-γ-methyleneadenosine 5’-triphosphate, AMPPCP). The results lead us to propose that stabilization of the P4 catalytic domain is key to the activation of CheA. We also propose a P1-P1’ dimer interface that remains dimeric throughout catalysis. Based on these findings we propose that changes in domain stability control the activation of CheA, a mechanism that may be important in the regulation of other kinases.

## Results

### Hydrogen exchange measurements of structure and dynamics of CheA within native-like complexes

HDX-MS provides insights into the structure and dynamics of proteins because the amide exchange rates depend on their environment (solvent exposure) and their role in the structure (eg involvement in backbone-backbone hydrogen bonds of the secondary structure). Application of this method to CheA in homogeneous, native-like signaling complexes provides insight into its physiologically relevant structure and what changes activate and inhibit the kinase. To achieve this, we optimized assembly conditions with excess CheW and CF to drive nearly all of the CheA into complexes (see Methods). These conditions are complementary to those of a previous study in which we used excess CheA and CheW to drive all of the CF into functional signaling complexes and measure its HDX properties (8), because HDX of each component of the complex must be measured in a separate set of experiments. The activity, stoichiometry, and stability of these complexes was measured to ensure that HDX experiments would reflect the properties of the CheA within functional complexes. Deuterium exchange was initiated with a spin column rather than dilution to ensure that complexes would not be dissociating during the time course of exchange. Triplicates of each HDX measurement were used to ensure reproducibility and to calculate the threshold for a significant change in HDX between states. Both the percent uptake and the exchange patterns were analyzed to identify key differences in CheA as it shifts between its kinase-on and kinase-off states. This approach yielded new insights into the stability and interactions of CheA within functional signaling complexes and how these change to modulate kinase activity.

### CheA in solution has unstable catalytic and regulatory domains

HDX-MS measurements on CheA in the absence of chemoreceptors and CheW should reveal domain stability and domain-domain interactions of CheA that contribute to its structure and function. We chose 3 µM CheA for our HDX experiments to (1) have predominantly dimeric CheA in solution, and (2) maximize the fraction of CheA bound in complexes (see next section). At this concentration, CheA alone in solution is about 75% dimeric, based on the K_d_=0.49 µM measured for *E. coli* CheA (18). Figure 2 illustrates the deuterium uptake of CheA relative to a fully exchanged control (CheA heat-denatured in D_2_O). The uptake time course is shown in Fig 2A for 46 representative peptides and for 4 of the timepoints collected. Data for all 91 peptides and 7 time points are shown in Fig S1. These peptides cover all but 69 residues (shown in black in Figure 2 and S1), providing HDX data for 89% of the 654 residues of CheA. The initial percent uptake (at 3 min) is represented on a structural model of the CheA dimer in Fig 2B. As expected, very rapid uptake (>90% in 3 min) is observed in the linkers between the P1 and P2 domains (L1) and between P2 and P3 (L2), consistent with prior studies indicating these domains are connected by flexible/disordered linkers (19, 20). Interestingly, the shorter L3 “hinge” connecting P3 and P4 also shows very rapid uptake, which suggests that, for CheA in solution, this hinge is not always in the β-strand conformation depicted in CheA models pdb *1B3Q* (21) and pdb *6S1K* (2).

**Figure 2.**
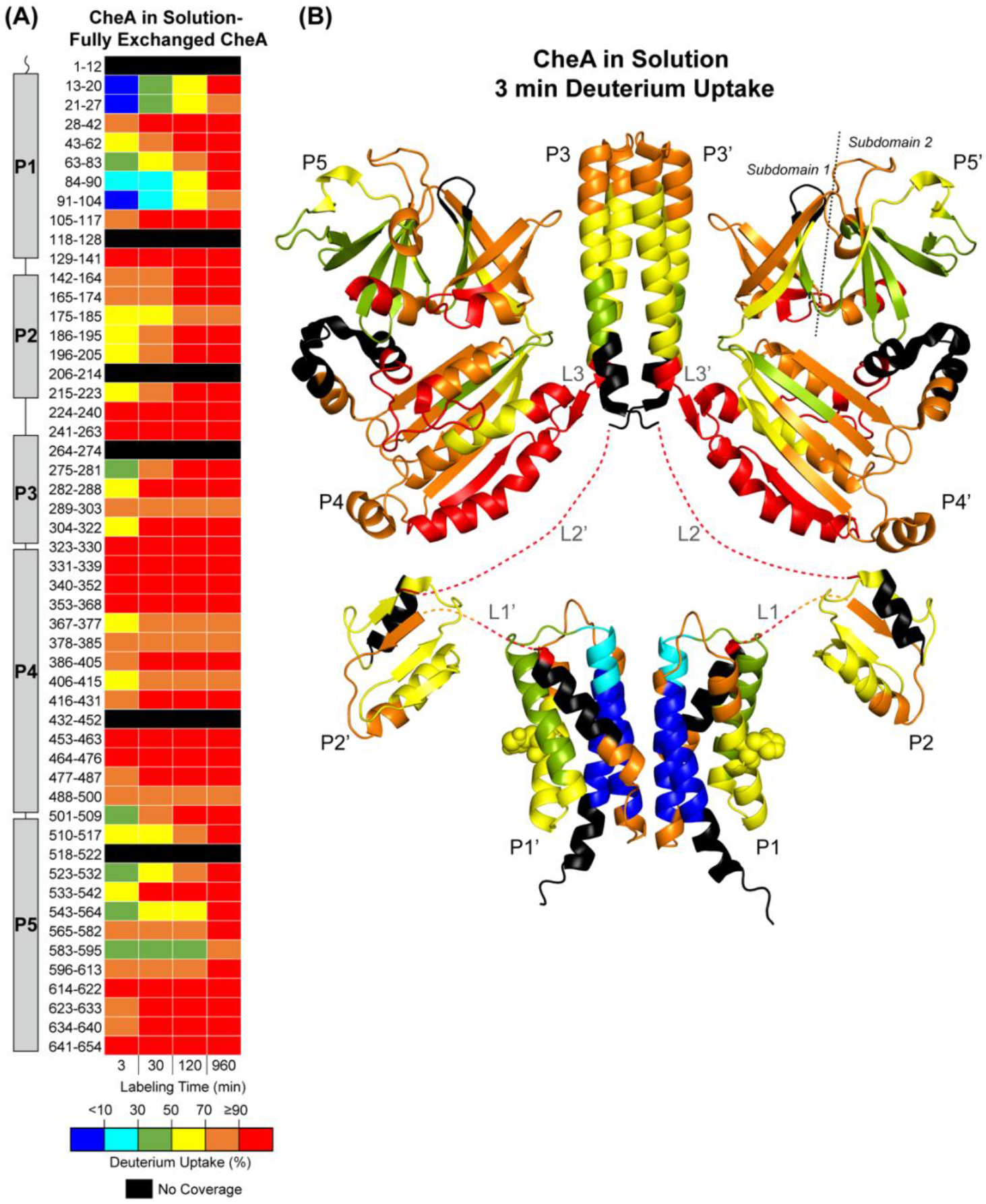
Deuterium uptake of CheA in the absence of signaling complexes. (*A*) Deuterium uptake versus time for selected peptides covering all five domains of CheA (cartoon at left) is represented in colors ranging from blue (<10% uptake) to red (≥ 90% uptake). Black indicates regions with no detected peptides. The uptake of each peptide was divided by the uptake of the same peptide in a fully exchanged CheA sample to determine percent uptake. (*B*) The uptake at 3 minutes for each peptide is represented on the structural model of a CheA dimer, consisting of pdb *6S1K* for the kinase core (P3P4P5) and pdb *2LP4* for P1 and P2, with dashed lines representing linkers that join P1 and P2 (L1) and P2 and P3 (L2), and the P1-P1’ interface modeled using HADDOCK. The phosphorylation site on P1 (His48) is shown in spheres.

Surprisingly, a significant fraction of the P4 (catalytic) and P5 (regulatory) domains also shows very rapid uptake (≥ 90% at 3 min). The stark difference between the HDX behavior of P4 and P5 vs. the other domains is summarized in Table 1, which gives the percent of each domain that undergoes rapid deuterium uptake (here defined as >70% uptake at 3 min, corresponding to red and orange colors in Fig. 2). Table 1 illustrates that P1, P2, and P3 are relatively stable domains for 3 μM CheA in solution, with 20-30% rapid uptake. In contrast, P4 and subdomain 1 of P5 exhibit about 70% rapid uptake. The rapid uptake of P4 suggests that this catalytic domain of CheA in solution is unstable. Since P4 is dramatically stabilized in kinase-on complexes, but not in partially kinase-off complexes (Table 1, and discussed below), we propose that kinase activity is regulated by stabilization of the catalytic P4 domain.

**Table 1.** Fraction of CheA domains with rapid deuterium uptake.

| Domain<sup>2</sup> | Sequence fraction with rapid<sup>1</sup> deuterium uptake |  |  |
| --- | --- | --- | --- |
| | Kinase-off<br>(3 μM CheA) | Partially kinase-off<br>(CheA in CF4E complexes) | Kinase-on<br>(CheA in CF4Q complexes) |
| P1 | 24% | 23% | 23% |
| P2 | 22% | 38% | 22% |
| P3 | 31% | 73% | 27% |
| P4 - catalytic | 74% | 70% | 28% |
| P5 – regulatory | 52% | 59% | 21% |
| P5 subdomain 1 | 65% | 65% | 0% |
| P5 subdomain 2 | 28% | 43% | 10% |
| L1 | 100% | 100% | 100% |
| L2 | 100% | 100% | 100% |
| L3 | 100% | 100% | 0% |

<sup>1</sup>Rapid defined as >70% deuterium uptake @ 3 min (red and orange colors in Figure 1). Reported sequence fractions are a lower limit, since peptides not detected were not counted as having rapid uptake.
<sup>2</sup>Domains are defined (39) as P1=1-131, P2=161-223, P3=262-325, P4=326-507, P5=508-654. P5 subdomain 1 = 510-530 and 596-634; P5 subdomain 2 = 531-595.

In contrast to the regions with rapid uptake, a striking feature of the HDX behavior of 3 μM CheA in solution is the very slow initial uptake observed on one face of the P1 domain. Very slow deuterium uptake (blue in Fig. 2) occurs only on two P1 helices: residues 13-27 of helix A (<10% at 3 min), and 84-104 of helix D (<16% at 3 min). These two helices of the five-helix bundle of *E. coli* CheA P1 are on the opposite side of the protein from the phosphorylation site (His48, shown in spacefill in Fig 2B). Since no other domains exhibit similar behavior, the slow uptake of helices A and D of P1 is likely due to a stable P1-P1’ dimer interface. This interpretation is consistent with prior evidence for P1 dimerization and with the fact that P1 deletion weakens the dimer affinity from K_d_ = 0.49 µM to 3 µM (18, 22). The slower uptake of the P1 dimer interface relative to the P3 dimer interface may be due to the fact that P1 forms a stable domain as a monomer, but P3 does not. The P1 domains were docked into a structural model for the dimer (discussed below), which is shown in all of the figures.

### Kinase core domains are stabilized in kinase-on complexes

To determine what changes in domain stability and interactions correlate with kinase activation and inhibition by chemotaxis receptors, we performed HDX-MS on CheA in native-like complexes, assembled on vesicles with the *E. coli* Asp receptor CF and coupling protein CheW. Complex assembly conditions were optimized to maximize the percentage of CheA bound, while maintaining high kinase activity (Fig. S2). Under limiting CheA conditions (3 µM CheA, 18 µM CheW, 30 µM CF, and 860 µM lipid), 80-90% of CheA is bound in functional complexes. CheA in these complexes exhibits high specific activity (23 s^-1^) in kinase-on complexes with CF4Q and CheW, and about half of that activity in complexes with CF4E and CheW (Fig. S3). The observed stoichiometry of 6 CF:0.5 CheA:3 CheW differs from the native 6 CF:1 CheA:2 CheW ratio, as expected for 90% binding of 3 µM CheA (predicts 6 CF:0.54 CheA), and binding of the homologous CheW to the sites that are missing CheA (predicts 6 CF:2.5 CheW). The kinase activity and stoichiometry do not change significantly over the 16 hour hydrogen exchange measurement period (Fig. S3). Thus, HDX of these samples reflects the properties of CheA in functional complexes. Although CF4E complexes under these conditions are not a perfect mimic of the kinase-off state, most likely due to crowded assembly on the vesicle surface (23), the differences in hydrogen exchange of CheA in CF4Q vs. CF4E complexes provide clear insights into the adaptation-related changes in structure and dynamics that activate and inhibit the kinase. Differences in deuterium uptake for each type of complex were determined by subtracting the centroid mass (intensity-weighted average mass) of each peptide for CheA in the complex from the centroid mass of that peptide for CheA in solution. Figure 3 illustrates the deuterium difference uptake for each type of complex, with both the time-course for representative peptides (top) and the 3 min difference uptake mapped onto the structural model of CheA (bottom). Blue shades indicate slower uptake in complexes, red shades indicate faster uptake in complexes, and gray indicates no difference, based on the calculated threshold of significance (see Methods). Difference uptake for all timepoints and all peptides of these complexes are shown in Fig S4.

**Figure 3.**
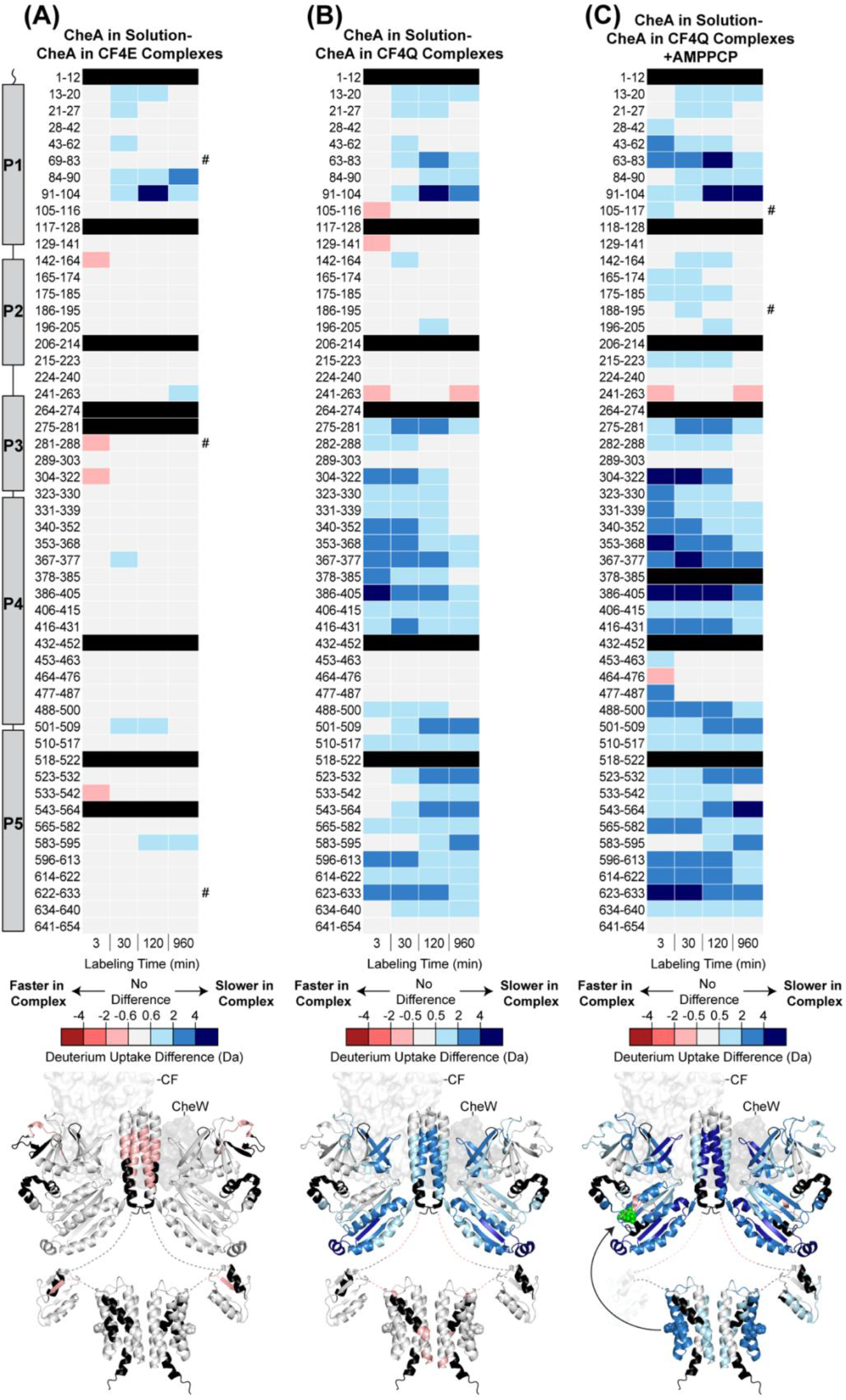
Deuterium uptake differences for CheA in signaling complexes vs in solution. Uptake differences for each peptide are calculated as uptake for 3 µM CheA in solution minus uptake for CheA in complexes, for *(A)* CF4E complexes, (*B)* CF4Q complexes, and *(C)* CF4Q complexes+AMPPCP. Difference uptake scale ranges from red indicating faster exchange of CheA in complexes to gray indicating no significant difference (see Methods for threshold calculation) to blue indicating slower exchange in complexes. Regions with no peptide coverage are shown in black. *Top*: time course of difference uptake for representative peptides. # indicates rows comparing similar but nonidentical peptides (see Figure S4 for the fully aligned difference uptake of all peptides). *Bottom*: uptake differences at 3 min are represented on the structural model of CheA (same model and orientation as shown in Fig 1), with the other elements of the complex shown in surface rendition: trimer of dimers of CF (white) and CheW (gray). AMPPCP is shown as green spheres in *(C, Bottom)*, and the black arrow indicates the likely interaction between P1 and P4 during catalysis.

Remarkably, there is very little change between the deuterium uptake of CheA in solution and that of CheA in complexes with CF4E and CheW (Fig 3A and Table 1). CheA is dimeric in complexes, and the array interactions will maintain a high local concentration of CheA, driving faster dimerization of any CheA that dissociates. Prevalence of the dimer should cause slower uptake at the P1 and P3 dimer interfaces. Consistent with this, reduced uptake is observed at longer times in complexes with CF4E for the P1 peptides suggested above to form the dimer interface (13-27 and 84-104). In contrast, P3 deuterium uptake is not slowed in CF4E complexes, and initial uptake is actually faster (Fig 3A, Fig S4, and Table 1), suggesting these complexes actively destabilize P3. Deuterium uptake is also unchanged in the P5 domain of CheA in CF4E complexes. This lack of change in the P5 domain is surprising, because within the functional complexes P5 forms an interface with CF and two interfaces with CheW. Thus, in CF4E complexes these interface interactions are too weak to stabilize P5 and too weak to decrease uptake at these interfaces. The third domain of the kinase core, P4, also shows no significant change in deuterium uptake. The lack of stabilization of the P4 catalytic domain of CheA in CF4E complexes likely accounts for the low kinase activity of CheA in its complexes with the demethylated state of the Asp receptor.

In contrast, CheA in kinase-on CF4Q complexes exhibits widespread slower uptake throughout the kinase core (P3,P4,P5), as shown in Fig 3B and Fig S4. For P4 and P5, the fraction with rapid exchange is dramatically reduced from 50-70% to 20-30%, suggesting the stability of these domains is now comparable to the P1, P2, and P3 domains (Table 1). In P5, the greatest stabilization is observed in subdomain 1, where CheA in solution exhibited rapid uptake (compare structures in Fig 3B with Fig 2B), suggesting strong interactions with CheW within core signaling units. Subdomain 2 of P5 also exhibits reduced uptake, especially at long times, consistent with stabilizing CheA/CheW interactions between core signaling units upon array formation in these samples assembled with the vesicle template methodology (8, 13). The incomplete exchange observed at 16 hours in P5 (Fig S1) is consistent with very stable interactions in the “baseplate” of the array between CheA, CheW, and the protein interaction region of the receptor, as shown in our previous HDX-MS study of CF in complexes (8). In contrast, P5 in CF4E complexes exhibits nearly complete exchange at 16 hours (Fig S1), consistent with our proposal that these protein interactions are weaker in the kinase-off state (8).The significant differences between the HDX of the kinase core of the CheA in CF4E complexes versus that of CF4Q complexes provides clear evidence that the stabilization of the kinase catalytic domain is required to fully activate CheA in chemoreceptor complexes.

Outside of the kinase core domains, there is no significant change in the initial uptake in P1, P2, and the linkers that connect them to the kinase core (Fig 3 bottom). At longer exchange times, reduced uptake is observed in P1(Fig 3B top), corresponding to stabilization of the P1 dimer interface (13-27 and 84-104) as well as another region to be discussed below. It is interesting that the P1 dimer interface is stabilized similarly in CF4Q and CF4E complexes, as expected due to stabilization of the CheA dimer in complexes, but only CF4Q stabilizes the P3 dimer interface. Finally, the hinge connector between P3 and P4 exhibits slower uptake in CF4Q complexes (Fig 3 and Table 1), indicating significant stabilization. Thus, inputs stabilizing the P4 catalytic domain in kinase-on complexes may include the stabilized P5, the stabilized P3 dimer interface, and the stabilized hinge connection between P3 and P4 (Fig 3B and Table 1).

### AMPPCP binding induces further stabilization that promotes transient interactions of P1 with P4

To further examine changes in CheA that occur during catalysis, we determined the effect of nucleotide binding on HDX of CheA in kinase-on complexes with CF4Q and CheW. We used a non-hydrolyzable ATP analog β-γ-methyleneadenosine 5’-triphosphate (AMPPCP), that is known to bind to CheA (24, 25). Measurement of AMPPCP inhibition of kinase activity of CF4Q complexes yielded an estimated inhibition constant (Ki) of 877 μM (Fig. S5). Samples of CF4Q complexes were prepared with 10 mM of AMPPCP (∼10X Ki) so that approximately 90% of CheA in complexes would have AMPPCP bound. HDX behavior of CheA in these complexes is shown on structural models that include the nucleotide; AMPPCP was positioned in the model by aligning pdb 1i58 (crystal structure of the isolated P4 domain of *Thermotoga maritima* CheA with AMPPCP) with the P4 domain of pdb 6S1K (RMSD=0.833). Fig 3C illustrates the two major effects of binding of AMPPCP on hydrogen exchange in 4Q complexes discussed below: (1) further stabilization of the kinase core, particularly the P4 catalytic domain, and (2) stabilization of a new face of P1.

Figure 4 shows the HDX behavior of the catalytic P4 domain in greater detail and illustrates where stabilization occurs in kinase-on complexes and upon nucleotide binding. P4 contains sequence motifs conserved among protein histidine kinases (PHK) that line the nucleotide binding pocket and include residues involved in ATP binding and Mg^2+^ coordination (21, 24, 26). Locations of the following conserved residues of these motifs are highlighted in Fig 4 (stick representations for sidechains, Cα spheres for Gly): His376 and Asn380 of the N box (on α2), Asp420 and Gly422 of the G1 box (following β3 and before α4), Phe455 and Phe459 of the F box (following α5), Gly470, Gly472, and Gly474 of the G2 Box (loop preceding α6) and Thr499 of the GT block (on β5). CheA in solution exhibits rapid uptake (>70% in 3 min, shown in red and orange in Fig 4A) for all of these regions containing these conserved residues, with the exception of His376 on α2, which shows slightly slower uptake. Crystal structures of *T. maritima* CheA P4 show the ATP lid (residues 461-471, between purple arrows in Fig 4B) to be poorly defined in the absence of nucleotides and to become more ordered upon nucleotide binding, including formation of a short α-helix in the P4-AMPPCP-Mg^2+^ complex (24). The ATP lid and flanking regions (residues 453-477) exhibit very rapid exchange in CheA alone (>90% exchange in 3min, red in Fig 4A) consistent with a highly flexible segment. The ATP binding pocket is significantly stabilized in kinase-on CF4Q complexes (Fig 4B): most segments containing conserved residues involved in ATP binding (Thr499) and Mg^2+^ coordination (His376 and Asn380) no longer exhibit rapid exchange in CF4Q kinase-on complexes, and exchange is even slower upon binding of AMPPCP (Fig 4C). An exception is the segment containing conserved ATP-binding residue Asp420, which exhibits rapid uptake (70-90%) in 4Q complexes and slows significantly to 30-50% uptake upon AMPPCP binding. Uptake remains very rapid (≥ 90%) in the F box – ATP lid – G2 box segment in CF4Q complexes (Fig 4B), with slower uptake only in the F box upon binding AMPPCP (Fig 4C). These results suggest no significant stabilization of the ATP lid, which likely continues to rapidly sample multiple conformations even with nucleotide bound. Notably, Bilwes et al suggested that the G2 box is not important for nucleotide binding affinity but may contribute to interactions with P1 during catalysis (26).

**Figure 4.**
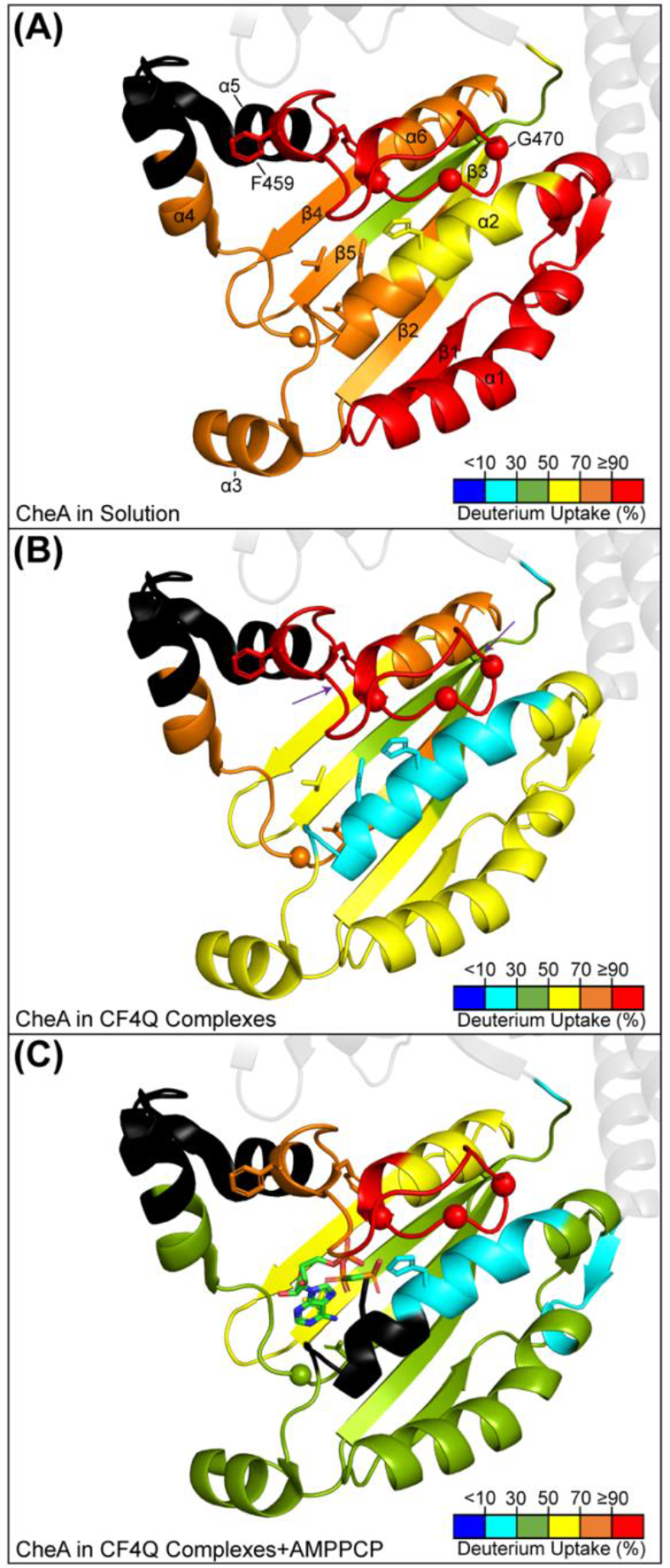
Catalytic domain deuterium uptake. (*A)* Percent uptake at 3 min relative to fully exchanged CheA of *(A)* 3 µM CheA in solution, (*B)* CheA in CF4Q complexes, and *(C)* CheA in CF4Q complexes with AMPPCP. Conserved residues are shown with stick sidechain representations or as spheres (glycine Cα’s). The segment between the purple arrows shown in *B* is the ATP lid (460-469). All structures are pdb 6S1K; AMPPCP (shown in stick representation in *(C)* was positioned by overlaying pdb *1i58* (P4 domain with AMPPCP).

The other important effect of AMPPCP on HDX of CheA in complexes is significant reduction in uptake in P1, in the area surrounding the phosphorylation site His48. Specifically, uptake is significantly reduced in both the His48-containing helix B (peptide 43-62) and the adjacent helix C (peptide 63-83), upon binding AMPPCP (Fig. 3C & Fig S4). Together these form a protein surface opposite the proposed P1 dimerization interface. Deuterium uptake is reduced in this region in CF4Q complexes at long exchange times (Fig 3B, top). Then, upon AMPPCP binding, there is even greater protection observed at both initial and long times (Fig 3C). It seems likely that nucleotide binding stabilizes the P4 domain sufficiently to enable the P1 surface formed by helix B and C to form a productive docking interaction and position His48 for *trans* phosphorylation (P1’ with P4). Note that slower uptake in the P1/P1’ interface helices (13-27 and 84-104) occurs whenever dimers are stabilized (CF4E, CF4Q, and CF4Q+AMPPCP complexes, Fig 3 top), but slower uptake of the P1 docking helices (43-83) occurs only upon stabilization of P4 (CF4Q ± AMPPCP, Fig 3B&C). In addition to the reduced uptake on both the dimerization and P4 binding surfaces of P1, we also observed widespread reduced uptake in P2 in the AMPPCP-bound state. Perhaps interactions of P1 with P4 also reduce the solvent exposure of P2.

### P1-P1’ dimer interface is maintained during signaling and catalysis

As discussed above, the very slow exchange of P1 helices A and D led us to propose that these helices form the P1/P1’ dimer interface. We generated a model of the P1-P1’ dimer using HADDOCK 2.4 (27), starting with the structure of *E. coli* P1 (residues 1-130) derived from pdb 2LP4. The HADDOCK approach for docking two proteins is to first define ‘active’ residues involved in the intermolecular interaction: residues that are both solvent-exposed and that are important in the binding, based on evidence such as chemical shift perturbation or mutagenesis. We used GETAREA (28) to identify solvent-exposed residues within the protected segments of helix A (residues 13-27) and helix D (residues 84-104) and designated these as active residues in the P1/P1’ interaction for HADDOCK docking with the default inputs. From the resulting clusters, we (arbitrarily) eliminated the antisymmetric dimer clusters, and chose the best scoring cluster, which generated a low Z-Score of -1.1.

The HDX results on CheA in complexes indicate that this dimeric P1-P1’ interaction persists regardless of the methylation state of CF or nucleotide binding to CheA. Furthermore, helices A and D exhibit slower uptake in all complexes, as expected since complexes stabilize the CheA dimer. This HDX behavior indicates that P1 remains dimeric when it transiently binds to P4 for phosphorylation during catalysis.

### Dimerization is not sufficient to activate CheA

We conducted additional HDX experiments at 0.5 µM CheA (50% dimer based on K_d_= 0.49 µM (18)) and 30 µM CheA (91% dimer), to compare to HDX of 3 µM CheA (75% dimer) and to HDX of CheA in kinase-on CF4Q complexes. The uptake difference between 0.5 µM and 3 µM CheA (Fig S6) indicates that the 25% increase in dimer fraction causes widespread reduction in HDX in both dimerization domains (P1 and P3) and throughout the kinase core (P3, P4, P5), with very little change observed in P2. Monomeric CheA is unstable, at least in the P3 and P4 domains, since the 0.5 µM CheA sample (50% monomer) exhibits very rapid uptake of nearly all the peptides in the P3 and P4 domains (red at 3 min in Fig S7, indicating >90% uptake). Dimerization clearly stabilizes CheA, but not enough to activate the kinase. First, we note that there is a negligible difference in dimer fraction of CheA in CF4E and CF4Q complexes, but a large difference in kinase activity. Based on the fraction bound and the K_d_=0.49 µM, we estimate that samples of CF4Q complexes have 94% dimer CheA (since 90% of 3 µM CheA is bound as dimers + 41% of 0.3 µM unbound CheA is dimeric) and samples of CF4E complexes have 91% dimer (since 80% of 3 µM CheA is bound as dimers + 50% of 0.6 µM unbound CheA is dimeric). Second, although dimerization stabilizes CheA (progressively reduced initial uptake for 0.5 to 3 to 30 µM CheA, Fig S7), P4 in particular retains rapid uptake at 3 min even for the 91% dimeric 30 µM sample (red and orange indicating >70% uptake, Fig S7), suggesting P4 remains unstable in CheA dimers. Finally, the difference uptake of 30 µM CheA vs CheA in CF4Q complexes (Fig S8), which have essentially the same 91-94% dimer fraction, demonstrates that incorporation into CF4Q complexes results in significant further stabilization of the kinase core. Thus the stabilization of the kinase core that occurs upon dimerization (Fig S6) is not enough to activate CheA; however, further stabilization of the kinase core in CF4Q complexes (Fig S8) correlates with kinase activation.

Bimodal HDX patterns provide further insight into CheA dimer properties because they were observed at both P1 and P3 dimer interfaces and can provide information about the timescale of exchange vs. dimer dissociation. Globular proteins typically exhibit uncorrelated exchange (EX2), where the isotopic distribution moves gradually to higher mass due to limited exchange occurring during multiple partial unfolding events, because the protein refolding rate is faster than the exchange rate (k_f_ >> k_exch_). Correlated exchange (EX1) occurs at the opposite extreme (k_f_ << k_exch_), because each unfolding event results in complete exchange for that peptide. With one exception (see below), EX1-like exchange is only observed in the dimer interface peptides of CheA, and thus seems likely to be due to dimer dissociation resulting in a long-lived open state of those regions. The rate of EX1 uptake is proportional to the rate of unfolding (in this case the dissociation rate), and the rate of EX2 uptake is proportional to the fraction unfolded (the stability). Fig S9 shows the mass spectra vs exchange time for P1 and P3 dimer interface peptides that exhibit clear EX1 and EX2 patterns. Percent uptake analysis ignores the underlying EX1 and EX2 processes, so these bimodal isotopic distribution patterns are analyzed by estimating the t_1/2_ of each process (8). Uptake is slow for the P1 dimer interface peptide (91-104) of 3 µM CheA in solution, with t_1/2_ of approximately 2 hours for both EX1 and EX2 (Fig S9C). In contrast, uptake is faster for the P3 peptide (304-322), with t_1/2_ of approximately 7 min for both EX1 and EX2 (Fig S9B). Relative to P3, the slower EX2 exchange of P1 suggests it is more stable and the slower EX1 suggests P1 dimers dissociate more slowly. Figure S10 further shows how the bimodal HDX patterns of the P1 and P3 interface peptides change for samples with different dimeric fractions: 50% dimer (0.5 µM CheA) to 75% dimer (3 µM CheA) to 91 % dimer (30 µM CheA). The observed. bimodal patterns reflect both the starting monomer/dimer fractions as well as dimer dissociation during HDX. Dimer dissociation is fast in P3, as demonstrated by the bimodal patterns of the P3 peptide: (1) the 50% dimer sample exhibits only the fast (monomer) HDX (complete at 3 min), and (2) the patterns for the 75% dimer and 91 % dimer samples are very similar. In contrast, the P1 peptide patterns exhibit (1) some slow (dimer) exchange for the 50% dimer sample, and (2) more distinct patterns for the 75% dimer and 91 % dimer samples. This further supports the conclusion that CheA dimers dissociate more slowly in the P1 domain than in the P3 domain.

The effects of complex formation on the EX1 and EX2 rates support the idea that CF4E destabilizes P3. With the exception of the P3 peptide in CF4E complexes, deuterium exchange spectra of signaling complexes exhibit clear stabilization for the P1 and P3 dimer interface peptides (Fig S9). The EX1 pattern disappears in spectra of the P3 peptide in 4Q complexes. Some EX1-like exchange remains in spectra for the P1 peptide in CF4E and CF4Q complexes, but the higher m/z envelope appears later and is no longer at the position of the fully exchanged sample. These observations suggest the open state closes more quickly, perhaps because any dissociated P1 or P3 dimer re-associates quickly within the signaling complexes. The increased t_1/2_ of EX2 reflects the stabilization of these dimerization domains. All complexes stabilize P1 (t_1/2_ of EX2 goes from 2 hours to >16 hours in CF4Q complexes). P3 only exhibits an increased t_1/2_ of its EX2 process in CF4Q complexes with AMPPCP; its slower overall uptake in CF4Q complexes is primarily due to elimination of the EX1 process. Interestingly, CF4E complexes increase the rate of EX1 and EX2 of the P3 interface peptide (t_1/2_ goes from about 7 min to 3 min), consistent with CF4E complexes destabilizing P3 and accelerating P3 dimer dissociation. Observation of EX1 in CF4E complexes suggests that the destabilized P3 cannot re-associate quickly and therefore exhibits a long-lived open state. Destabilization of P3 by interactions with CF4E may be the mechanism by which receptor methylation state modulates kinase activity.

Finally, the kinase core of monomeric CheA may be partially disordered, with a long-lived open state. The only other EX1 processes observed for CheA in this study occur in several peptides of P5 (523-532, 543-564, 553-564, 583-595) of the 50% monomeric 0.5 µM CheA sample (Fig. S11). These are the slowest exchanging peptides of the P3, P4, and P5 domains in this sample (see Fig S7), which makes it possible to observe EX1. Other peptides of the kinase core may also undergo EX1 HDX that would not be detected due to their rapid exchange. EX1 is not detected in the P1 and P2 peptides (except the P1 interface peptide, as discussed above), suggesting these are stable domains in monomeric CheA. Overall, it appears that the kinase core domains are unstable and partially disordered in CheA monomers.

## Discussion

Based on the hydrogen exchange properties of CheA, we propose that chemoreceptor signaling complexes control its kinase activity via stabilization/destabilization of its catalytic domain P4, as depicted in Figure 5. This proposed mechanism is based on the observation that for states of CheA with low kinase activity (CheA in solution and in CF4E complexes), P4 exhibits rapid deuterium uptake, suggesting low stability of this domain. Furthermore, activation of CheA in CF4Q complexes correlates with significantly slower deuterium uptake in P4, suggesting stabilization of the catalytic domain activates the kinase. In Figure 5 stable domains of CheA are shown in blue, such as P1 and P2 that exhibit a low (20-30%) fraction with rapid deuterium uptake for all states (Table 1). Unstable domains are shown in red, such as P4 and P5 that exhibit a high (50-70%) fraction with rapid uptake for CheA in solution, even for 90% dimeric CheA (Fig S7). By these criteria, CheA in solution (Fig 5A) has a stable P3 only in dimeric CheA (3 and 30 µM CheA with 75-90% dimers, see Figure S7). Upon assembly of CheA into kinase-activating CF4Q complexes (Fig 5C), deuterium uptake is reduced throughout the kinase core domains (Fig 3B) and the P4 and P5 domains become stabilized (20-30% rapid uptake, Table 1). Finally, nucleotide binding to P4 (Fig 5D) causes reduced deuterium uptake in both P4 and on the P1 surface that bears the His48 phosphorylation site, likely by promoting the P1/P4 interactions needed for catalysis. This result is consistent with the ordered sequential binding mechanism suggested by recent enzyme kinetics measurements, in which ATP must first bind to P4 to create a docking surface for P1 to bind for phosphoryl transfer (25).

**Figure 5.**
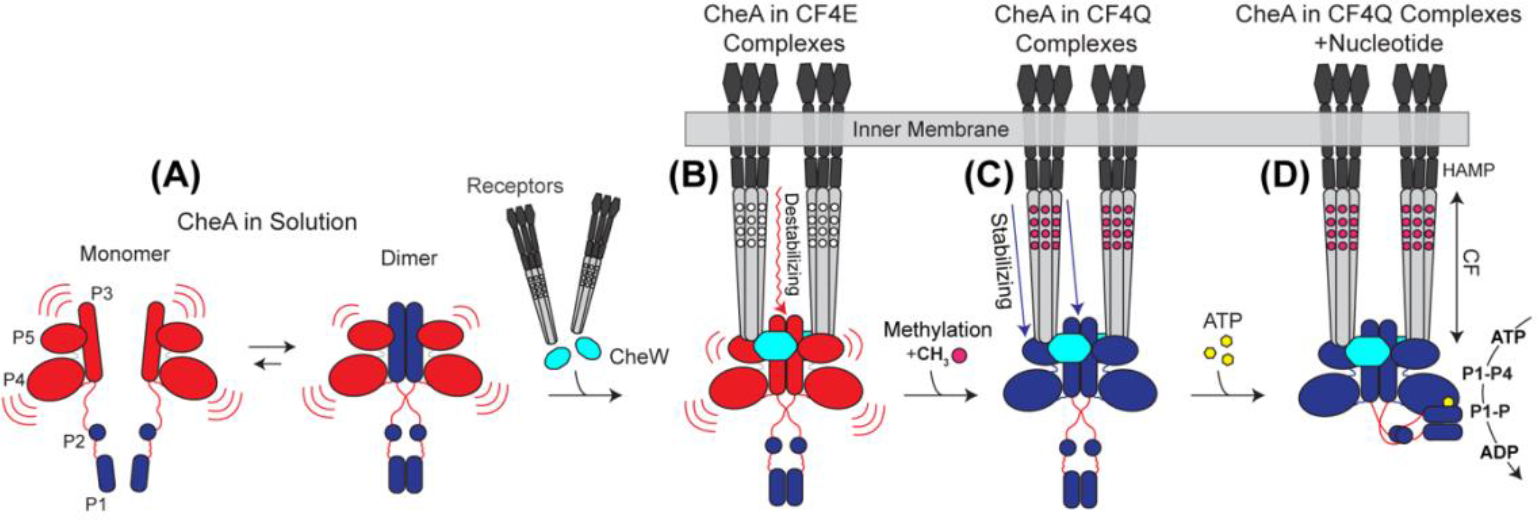
Model for control of CheA kinase activity by chemotaxis signaling complexes. Note that HDX experiments were performed on CF complexes lacking the black segments of the receptor shown in the model. (*A*) CheA in solution: Monomeric CheA has an unstable kinase core (P3,P4,and P5 shown in red) connected by flexible linkers to the stable (blue) P1 and P2 domains; dimerization stabilizes primarily the P3 domain. The instability of the P4 catalytic domain accounts for the low kinase activity of CheA in solution. *(B)* Assembly into complexes with unmethylated receptors (CF4E in our experiments) and CheW (cyan) destabilizes P3: the kinase core is not stabilized, and kinase activity is low. *(C)* In contrast, complexation with the methylated receptor (magenta circles represent methylated sites, mimicked by CF4Q) stabilizes the kinase core, presumably through stabilizing interactions with P3 and P5, which in turn stabilize P4. *(D)* ATP binding (yellow hexagon) further stabilizes the kinase core to promote interactions between P4 and a dimeric P1-P1’ for phosphoryl transfer to His48 on P1’.

HDX rates are affected by both solvent accessibility and protein structural fluctuations. Covalent labeling with diethylpyrocarbonate is a complementary approach, because the slower timescale of the labeling makes it insensitive to structural fluctuations (29). Our recent covalent labeling study of CheA in solution and in CF4Q complexes (30) provides insights into the causes of reduced HDX in P4 and P5 in CF4Q complexes. Several sites in P4 and P5 show decreased HDX rates but no change in covalent labeling (Table S1), suggesting these sites experience less structural fluctuations in CF4Q complexes. This is consistent with our interpretation that domains P4 and P5 are stabilized in CF4Q complexes.

Interestingly, domain stabilization is thought to be important in the activation mechanism of the EnvZ His kinase: HDX-MS data suggest that high osmolarity enhances kinase activity by stabilizing the partially disordered domain containing the His phosphorylation site of EnvZ (31). In contrast, our HDX data on CheA show no evidence for a partially disordered P1 domain in CheA dimers or complexes (eg see Figs S9-S10 and S12, in which bimodal envelopes are all consistent with dimer dissociation, as discussed above), and its H48 phosphorylation site is not part of the His-Asp/Glu dyad observed in EnvZ and some other His kinases (31). CheA activation instead involves stabilization of its catalytic P4 domain.

Hydrogen exchange measurements on CheA in partially kinase-off CF4E complexes reveal a possible mechanism for adaptational control of the kinase. These complexes fail to stabilize CheA (no significant reduction in deuterium uptake, Fig 3A). Instead CF4E complexes (Fig 5B) apparently destabilize P3, as seen in increased overall deuterium uptake in P3 (Figs 3A and S4) and faster rates of both EX1 and EX2 hydrogen exchange in P3 (Fig S9). We suggest there are destabilizing interactions between the unmethylated receptor and P3, and this P3 destabilization interferes with stabilizing interactions with P5 in the complex. This is consistent with our previous proposal that there are weaker binding interactions between the receptor and CheA in kinase-off complexes (8).

The observed instability of the *E. coli* CheA kinase core may not be a shared feature with the other well-studied *Thermotoga maritima* CheA. *T. maritima* and *E. coli* exhibit opposite regulation of CheA: *T. maritima* CheA is active in solution and inhibited by chemoreceptor complexes; *E. coli* CheA is inactive in solution and activated by chemoreceptor complexes (22, 32, 33). Also, ATP and a fluorescent nucleotide analog bind to *T. maritima* CheA over 10-fold more tightly than to *E. coli* CheA (24, 34). It is notable that most structural studies of the kinase core domains have been performed on *T. maritima* CheA, including x-ray crystal structures of P4 (24) and P3P4P5 (21). In contrast, structural studies of *E. coli* CheA domains are typically limited to P1 and/or P2 (35–39). These previous studies are consistent with the idea that *E. coli* CheA P1 and P2 are stable domains, but *E. coli* P3P4P5 is not stable until incorporated into kinase-on signaling complexes.

In addition to leading to the proposed role of stabilization/destabilization of *E. coli* CheA domains P4 and P3 in kinase activation and adaptation, HDX results provide insights into the P1/P1’ dimer. P1 is known to contribute to the dimerization of CheA, as deletion of P1 increases K_d_ from 0.49 µM to 3.1 µM (18). Slow exchange of some residues within helices A (residues 4-30) and D (residues 84-107) was previously observed in an NMR study of isolated (presumably monomeric) P1 (40), suggesting these helices are especially stable and/or buried, even in the monomer. Dimerization causes reduced deuterium uptake throughout P1 (Fig S6), suggesting widespread stabilization. What identifies the A/D surface as the dimer interface is that it shows reduced deuterium uptake in all complexes, active or inactive, which makes it unlikely that protection is due to a non-productive interaction with P4. Instead, the reduced uptake of helices A and D in all complexes is likely due to increased dimer fraction and/or faster dimer reassociation rates in the complexes. Therefore, residues in helices A and D that are both solvent-exposed and very slow exchanging were used as docking constraints in HADDOCK to yield a structural model for the P1 dimer. As shown in Fig S13, the proposed *E. coli* P1 dimer model differs from a crystallographic dimer of *T. maritima* CheA P1 (pdb 1tqg), which places primarily helices A and B at the dimer interface (22). The dimer interface formed by the A and D helices in our *E. coli* P1 dimer model is mostly adjacent to the sites on the A and B helices suggested by a mutagenesis study to be important for P1/P4 interactions (41). Thus, the HDX data suggest that P1 dimerization does not interfere with phosphorylation and instead this domain remains dimerized throughout the catalytic cycle. This evidence that P1 dimers do not dissociate during phosphorylation is consistent with the previous suggestion that *E. coli* CheA in solution only phosphorylates one P1 at a time (half-of-sites reactivity), based on 5-fold higher rates of autophosphorylation measured in heterodimers containing a single P3P4 compared to homodimers of intact CheA (42).

The hydrogen exchange properties of the linkers between CheA domains provide further insights into signaling-related changes in domain interactions. The long linkers L1 (134-159) and L2 (229-261) that connect P1 and P2 to the kinase core have very rapid (≥ 90%) or rapid (>70%) uptake in all states (Table 1, Fig 2 and Fig S1). This suggests that P1 and P2 retain mobility and experience only transient interactions with other domains. In contrast, signaling-related changes are observed in the deuterium uptake of the short linkers L3 and L4, both of which are thought to be important in signaling (43–45). As highlighted in Table 1, L3 (322-327) undergoes very rapid exchange in kinase-off states (>90% in 3 min, Fig 2), which is slowed significantly in kinase-on states (to 59% in CF4Q and then 26% upon addition of AMPPCP, Fig S1). Although L4 (506-509) has slower exchange (49% at 3 min in kinase-off states, Fig S1), it also exhibits reduced deuterium uptake in kinase-on states, particularly at longer times (Fig 3 and Fig S4). These short linkers flanking P4 control its position, for instance to allow a P4 “dipped” conformation observed by molecular dynamics (46), which is suggested to correspond to the kinase-on state based on mutagenesis studies (47). We suggest that P4 samples multiple positions in the kinase-off state (e.g., undipped and dipped) leading to very rapid HDX of L3. Upon kinase activation, L3 adopts a single conformation that positions P4 (perhaps in the dipped conformation) for productive interactions with the P1/P1’ dimer and phosphoryl transfer to P1’. In this model, P1 and P2 would localize primarily below the complex in the kinase-off state; upon kinase activation, these domains spend less time in this position due to transient interactions with P4, as shown in Fig 5D. This could account for the smaller “keel” of P1/P2 electron density observed beneath the complex in cryo-electron tomography studies of complexes of 4Q receptors relative to 4E (48). In this view, P1 and P2 remain mobile in both kinase-off and kinase-on complexes, but in the kinase-on state they spend more time in a compact conformation interacting with P4 (Fig 5D). This more compact conformation may protect L1 and L2 from proteolysis, and thus account for the observation of reduced proteolysis of CheA in 4Q complexes (48). This contrasts with the observation that P1 is released upon AMPPCP binding, resulting in a less compact conformation; however this effect was primarily observed for *T. maritima* CheA, with a much smaller change observed for *E. coli* CheA (49).

In principle, the CheA kinase can be activated by promoting productive P1/P4 interactions and can be inhibited by promoting nonproductive P1/P4 interactions. The P4 side of P1/P4 interfaces could not be identified in this hydrogen exchange study, due to the properties of P4 – widespread rapid deuterium uptake in kinase-off states and widespread stabilization in kinase-on states. P1 has more favorable HDX properties that revealed two interaction interfaces. (1) Similarly reduced deuterium uptake of helices A and D in both kinase-on and kinase-off complexes is indicative of the P1/P1’ dimer interface that persists through catalysis. (2) Reduced uptake in helices B and C in kinase-on complexes, particularly upon binding of AMPPCP to P4, is indicative of the productive P1/P4 interaction for phosphorylation of His48 in helix B. We saw no protection of P1 that occurred only in the partially kinase-off complex with CF4E, and thus no evidence of the non-productive P1/P4 interaction observed between P1 and P3P4 of *T. maritima* CheA, which has been proposed to sequester P1 in the kinase-off state (50). It is possible that this nonproductive P1/P4 interaction is specific to *T. maritima* CheA, or that it occurs at the P1 dimerization interface when P1 is a separated domain at concentrations too low to dimerize.

Finally, our results are consistent with an ordered sequential binding mechanism for CheA, in which ATP binding to P4 promotes subsequent binding of P1. Recent kinetic analysis of *E. coli* CheA P3P4P5 phosphorylation of P1 in signaling complexes and binding of P3P4P5 to P1 both support this mechanism (25). Consistent with this proposal, significantly more protection of P1 helices B and C is observed in the presence of AMPPCP, suggesting nucleotide binding is needed to promote productive P1 interactions with P4. However, the ATP lid is not ordered in the nucleotide-bound complex; it still undergoes very rapid deuterium uptake (Fig 4C). In light of their evidence for an ordered sequential binding mechanism, Hazelbauer and coworkers propose that receptor complexes modulate kinase activity by controlling ATP binding (25). In another kinetic study of CheA, these authors reported that signaling complexes primarily modulate k_cat_, achieving activation with a ∼100-fold increase upon incorporation into signaling complexes and inhibition with a ∼40-fold decrease upon ligand binding (10). Our proposal that kinase activity is modulated by stabilization/destabilization of the catalytic P4 domain is consistent with both of these studies. Stabilization of P4 as a mechanism of kinase activation should promote binding of ATP, binding of P1, and catalysis, thus modulating multiple kinetic parameters. Such a powerful control mechanism is likely used to modulate kinase activity in other systems, particularly within complexes like the chemotaxis signaling complex that may protect the unstable domain of the kinase-off state from proteolytic degradation.

## Materials and Methods

Sample preparation, kinase assays, and HDX data acquisition and analysis are fully described in the ***SI Appendix, Materials and Methods***.

### Protein purification

All proteins used in this study were *E. coli* chemotaxis proteins. The His-tagged Asp receptor cytoplasmic fragment (CF), and TEV-cleavable, His-tagged CheA, CheW, and CheY were purified according to previously established protocols (8, 14).

### Vesicle mediated assembly

Complexes were assembled by combining CheA, CheW, vesicles, and CF4Q or CF4E, followed by overnight incubation at 25°C. Kinase and sedimentation assays were used to assess sample functionality and the fraction of each protein bound (8).

### Sample preparation for HDX-MS

Functional signaling complexes were prepared with 3 µM CheA, 18 µM CheW, 30 µM CF (CF4Q or CF4E), and 860 µM lipid. Samples were transferred into deuterated buffer using desalting columns and then incubated at pD 7.5, 25°C before quenching at pH 1.6, 0°C and flash-freezing.

### HDX-MS data acquisition and analysis

MS data were acquired in the UMass Amherst mass spectrometry core facility on a Waters SYNAPT G2Si mass spectrometer with IMS. Three replicates were assayed for every state and time point (except only two replicates for the 30 µM CheA sample). Data analysis was performed using Protein Lynx Global Server (PLGS) version 3.0.1 for peptide identification and DynamX version 3.0 for determination of deuterium uptake.

## Acknowledgments

Mass spectrometry data were obtained at the University of Massachusetts Mass Spectrometry Facility with support from the Institute for Applied Life Sciences. We thank Sandy Parkinson, Richard Vachet, and Katie Wahlbeck for helpful comments on the manuscript. This work was supported by National Institutes of Health Grant R01-GM120195 (to L.K.T.).

