## Supplemental Information for "Bacterial chemoreceptor signaling complexes control kinase activity by stabilizing the catalytic domain of CheA"

[*]Corresponding Author:

**This PDF file includes:**

Supplementary text
Figures S1 to S13
Table S1
SI References

### Supplementary Information Text

#### Materials and Methods

**Protein purification.** All proteins used in this study were *E. coli* chemotaxis proteins. The His-tagged Asp receptor cytoplasmic fragment (CF) was purified according to previously established protocols (1, 2). The Asp receptor has four sites where methylation and demethylation of glutamate residues mediates adaptation. In the wild type receptor, two of these sites are initially encoded as glutamine (Gln295, Glu302, Gln309, Glu491), which poises the receptor in the half-methylated state since glutamine mimics the activity effects of the glutamyl methyl ester (3). The fully unmethylated and methylated-mimic His-tagged CF were expressed using the plasmids pHTCF4Q(amp$^r$) and pHTCF4E(amp$^r$) (4). They were co-transformed with pCF430 (tet$^r$) encoding lacI$^q$ plasmid into BL21(DE3) cells. Plasmids pHisTEV-CheA (kan$^r$), pHisTEV-CheW (kan$^r$), and pHisTEV-CheY (kan$^r$) encoding TEV-cleavable, His-tagged CheA , CheW, and CheY, respectively, were expressed in BL21(DE3) cells and also purified as previously described (2). Protein concentrations were determined using BCA assays (CF) or absorbance at 280 nm with extinction coefficients (5, 6) of 25,000 M cm$^{-1}$ (CheA), 5120 M cm$^{-1}$ (CheW), and 10,700 M cm$^{-1}$ (CheY).

**Lipid vesicle preparation.** Large unilamellar vesicles for assembly of functional complexes of CF, CheA and CheW were prepared as previously described (2), using DOGS-NTA-$Ni^{2+}$ (1,2-dioleoyl-sn-glycero-3-[(N-(5-amino-1-carboxypentyl) iminodiacetic acid)succinyl] (nickel salt) and DOPC (1,2-dioleoyl-sn-glycero-3-phosphocholine) obtained from Avanti Polar Lipids Inc, AL. Briefly, DOGS-NTA- $Ni^{2+}$ and DOPC were mixed in a molar ratio of 1:1.5, chloroform was removed, and lipids were hydrated in kinase buffer (50 mM $K_xH_xPO_4$, 75 mM KCl, 5 mM MgCl2, pH 7.5). After hydration and five freeze thaw cycles, vesicles were extruded through a 100 nm pore polycarbonate membrane to form unilamellar vesicles. Final total concentrations of 860 µM lipid were used in complex assembly to provide sufficient vesicle surface area to assemble 30 µM CF in functional signaling complexes (8).

**Vesicle mediated assembly.** Complexes were assembled by combining the proteins and vesicles in kinase buffer, in the following order: CheA, CheW, vesicles, and CF4Q or CF4E. After overnight incubation at 25˚C, an enzyme-coupled kinase assay was used to determine the functionality of the complex and a sedimentation assay was used to determine the fraction of each protein bound to the vesicles, as previously described (1).

Limiting CheA conditions were needed for measurements of the HDX properties of CheA in complexes. Assembly conditions were optimized to achieve active complexes with at least 80% of the CheA bound. The native stoichiometry of 6 receptor:1 CheA predicts a maximum of 5 µM CheA will assemble into active complexes with 30 µM CF. We varied the CheA concentration, keeping a molar ratio of 1 CheA:2 CheW, and found high activity with 3 µM CheA. For fixed concentrations of 3 µM CheA, 30 µM CF4Q, and 860 µM lipid, the CheW concentration was varied (Fig S2) and the maximal activity and bound fraction of CheA was observed for 18 µM CheW. Hence, all samples of signaling complexes for MS experiments were prepared with these conditions.

**Kinase assays.** The assay monitored the decrease in absorbance at 340 nm due to oxidation of NADH, which was coupled to ATP consumption. For each measurement, 2 µL of complex was mixed with 198 µL of CheY mix (55 µM CheY, 20 units of PK/LDH enzyme, 4 mM ATP, 2 mM phosphoenolpyruvate, and 250 µM NADH) in kinase buffer, and absorbance was measured for 72 s. Kinase activity was calculated by subtracting the slope of the absorbance change at 340 nm of a CheY-only background control from the slope measured for the complex. Specific kinase activity per total CheA was calculated as follows: [adjusted slope as abs/s) X (M NADH/6220 abs)] / [M CheA total) X (2 µL / 200 µL)]. Initially, to check the functionality of purified proteins, complexes were assembled with excess CheA and CheW (12 µM CheA, 24 µM CheW, 30 µM CF4Q).

**Sample preparation for HDX-MS.** Functional signaling complexes were prepared with 3 µM CheA, 18 µM CheW, 30 µM CF (CF4Q or CF4E), and 860 µM lipid in Kinase buffer and incubated overnight at 25˚C. The total volume of each sample was 1 mL. Before initiating exchange, kinase activity was measured and sedimentation assays were performed to measure the fraction bound of CheA, CheW and CF. Deuterium exchange was performed using 99.9% $D_2O$ (Cambridge Isotopes) and desalting columns (2 mL Zeba desalting column, ThermoFisher). Columns were pre-equilibrated with deuterated kinase buffer (50 mM $K_xH_xPO_4$, 75 mM KCl, and 5 mM MgCl2 at pD 7.5 corresponding to a pH reading of 7.1) at 25°C as follows: spin column at 2000 rpm for 2 min in a tabletop centrifuge (Beckman Coulter Allegra R Tabletop Centrifuge), discard the flow-through, add 1 mL deuterated buffer to the top of the column, spin again and repeat until the column has been exchanged with a total of 4-5 ml of deuterated buffer. The verified active complex (1 ml) was added to the pre-equilibrated column, followed by centrifugation at 2000 rpm for 2 min at 25°C to initiate deuterium exchange. The eluted solution of complexes was incubated in a 25°C water bath and aliquots were removed after exchange for 3 min, 7 min, 15 min, 30 min, 60 min, 120 min, and 16 hr (note that the aliquot collected immediately after the 2 min spin + 1 min deceleration represents 3 min of exchange). For each time point of exchange, 22.5 µL was transferred into a pre-chilled Eppendorf tube containing 22.5 µL quench buffer (1% formic acid, 20% w/v glycerol, 1 M GuHCl, pH 1.6) in a 0°C ice-water bath. Immediately after quenching, each sample was vortexed, flash-frozen in liquid nitrogen, and stored at -80°C until the MS experiment.

Free CheA samples were prepared in a similar manner, omitting the vesicles and other proteins; undeuterated CheA samples were prepared by omitting the exchange steps.

To prepare the fully exchanged control, 3 μM CheA was exchanged into deuterated kinase buffer using a desalting column, following the same protocol as above. Exchanged CheA was heat denatured in a thermocycler (BioRad) by heating to 95°C for 30 min; then it was cooled to 25°C at a rate of -3°C/min, and incubated in a 25°C water bath for 20 h. Then quenching, freezing, and MS steps were conducted in the same manner as for all of the other samples, so that this fully exchanged control would undergo the same extent of back-exchange. Using this fully exchanged control, we calculated back-exchange percentage as follows (7): (1) the percent of the theoretical maximum uptake was calculated for each analyzed peptide as 100*(measured uptake/ number of exchangeable amides of backbone in peptide). (2) Percent uptake was then divided by 0.91 to account for an estimated 10% dilution during $D_2O$ exchange via spin column, and then subtracted from 100 to yield the back exchange level. The average back-exchange percentage was 18% (range of 3-36%).

The nucleotide-bound state of CheA in complexes was prepared by addition of the inhibitor β,γ-methyleneadenosine 5'-triphosphate (AMPPCP; Sigma) to CF4Q complexes. To estimate Ki, kinase activity of CheA in CF4Q complexes was measured as a function of [ATP], in the presence of several concentrations of AMPPCP. The CheY mixture included [AMPPCP], for final concentrations of 0, 0.8, 1.6, or 3.2 mM in the assay. Global fits of competitive inhibition were performed with OriginPro to determine a Ki of 877μM. We chose 10 mM AMPPCP to add to HDX samples, which predicts that 92% of CheA will be bound to AMPPCP if Kd = Ki = 877 μM. CheA in CF4Q complexes samples was prepared as described above. After verifying activity and complex formation, the sample was incubated with 10 mM AMPPCP for 5 min at 25°C. The desalting column for exchange was prepared by exchanging it 3 times with deuterated kinase buffer, followed by a final exchange with deuterated kinase buffer supplemented with 10 mM AMPPCP. Samples were exchanged, quenched, and stored as described above.

**HPLC column preparation and maintenance.** For peptide separation in mass spectrometry experiments, a C18 reverse phase HPLC column (2.1 mm x 5 cm, Higgins Analytical) was used. Before every set of MS experiments, the column was connected to an LC pump (Agilent 1100 G1312A) and cleaned with Buffer B (0.1% formic acid in acetonitrile) by running at a flow rate of 200 μL/min overnight. At the end of the set of MS experiments, the column was stored with 100% Buffer B. Every two months the column was subjected to deep cleaning by flushing at 200 μL/min with 20 column volumes of water, 20 column volumes of acetonitrile, 5 column volumes of isopropanol, 20 column volumes of heptane, 5 column volumes of isopropanol, and then 20 column volumes of acetonitrile.

**HDX-MS data acquisition.** MS data were acquired in the UMass Amherst mass spectrometry core facility (RRID: SCR_019063) on a Waters SYNAPT G2Si mass spectrometer with IMS. Before running samples, the mass spectrometer was calibrated in positive ion mode using sodium iodide. All samples were acquired in positive ion mode with ion mobility separation in the mass range 100-2000 m/z. A 100 fmol/μL solution of leucine enkephalin ([M+H]+ 556.2771 m/z) was continuously infused at 2 μL/min for lock mass correction. Valves and tubing were cleaned thoroughly with methanol, water and 50% isopropanol before each use. Prior to every MS run, the HPLC column was washed with 95% of Buffer B (0.1% formic acid in acetonitrile) and equilibrated with 95 % Buffer A (0.1% formic acid in water). We followed the method used by Li et al. to minimize back-exchange (1). Briefly, the analytical C18 column, connecting polyetheretherketone (PEEK) tubing and valves were placed in an icebox to maintain a constant temperature of 0°C. Frozen, deuterium-exchanged, and quenched complexes were thawed for 1 min in a 0°C ice-water bath and digested for 1.5 min with 40 μM pepsin (Sigma Aldrich). Then the sample was immediately injected via the injection valve onto the chilled column. Peptides were eluted at 100 μL/min directly into the mass spectrometer. 5% Buffer B was flowed through the column for 3 min for desalting followed by a 5-50% Buffer B gradient over 12 min for peptide separation. After data acquisition the column was washed with 95% Buffer B for 27 min to elute the remaining peptides, followed by 5% buffer B to re-equilibrate the column. The initial 3 min desalting and 27 min post-gradient elution and wash steps were diverted to waste to minimize ion source contamination. A blank injection (5% Buffer B) was performed after the column wash to ensure no peptide carryover before injecting the next sample. All deuterium-exchanged samples were run in random order and three replicates were assayed for every state and time point (except only two replicates for the 30 µM CheA sample). Each set of MS experiments also included an undeuterated CheA sample run with collision-induced dissociation in $MS^E$ mode to confirm peptide identification, and a fully-exchanged control sample to measure the back exchange.

**HDX data analysis.** Peptide identification was conducted using Protein Lynx Global Server (PLGS) version 3.0.1 software from Waters. Data acquired from undeuterated samples were searched against sequence database of proteins present in the complexes and pepsin. Processing parameters include 556.2771 Da/e as lock mass for charge 1, lock mass window of 0.25 Da, and non-specific primary digest reagent.

PLGS results were imported to Waters, DynamX version 3.0 software for determination of deuterium uptake of samples. PLGS filtering parameters were as follows: minimum signal intensity of 20000, minimum products per amino acid of 0.3, maximum sequence length of 25 amino acid and retention time RSD of 0.5%. After the automated processing by DynamX, every spectrum was manually checked for correct retention time, m/z and charge state assignments. Peptides with inconsistent data (one of the triplicates deviates significantly) or bad spectra were removed from the analysis. Then uptake data was exported to Excel for further analysis. Uptake data were corrected for back exchange using data from the fully-exchanged control, to generate percent uptake heat maps. Differences in uptake between two states were calculated to generate difference uptake heat maps. The following approach was used to determine the threshold (in Da) for a significant difference in uptake between two different states. The uptake standard deviation (between the triplicate measurements) for all time points of all peptides was averaged for each state. These two average standard deviations were used to calculate the standard error difference, which was multiplied by the Student's t-table value for a 95% confidence interval and four degrees of freedom to determine the threshold for a significant uptake difference. Uptake percentage and differences in uptake were mapped onto a structural model consisting of pdb 6S1K (P3, P4, and P5 domains) and pdb 2LP4 (P1 and P2 domains).

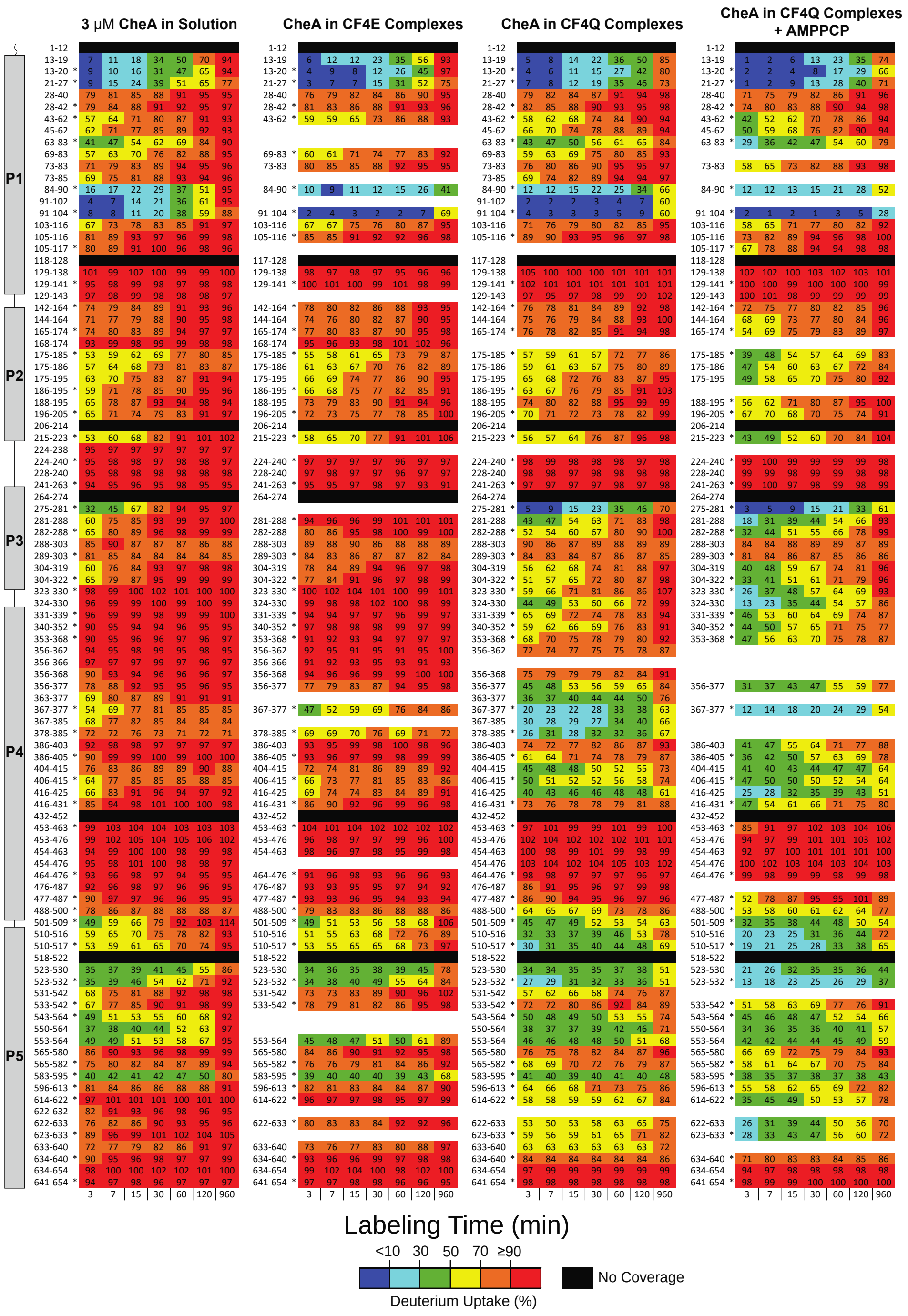


**Fig. S1.** Deuterium uptake percentage vs time for all peptides and time points in each state of CheA. Uptake percentages of CheA peptides were calculated by dividing uptake in that state by uptake of a fully exchanged CheA sample. Uptake scale ranges from blue representing low uptake (<10%) to red representing high uptake (≥90%). Asterisks indicates peptides chosen for representing uptake and uptake differences on CheA structures in Figures 2-4. Black bars indicate regions with no coverage; white gaps indicate missing peptide, but other peptides provide coverage.

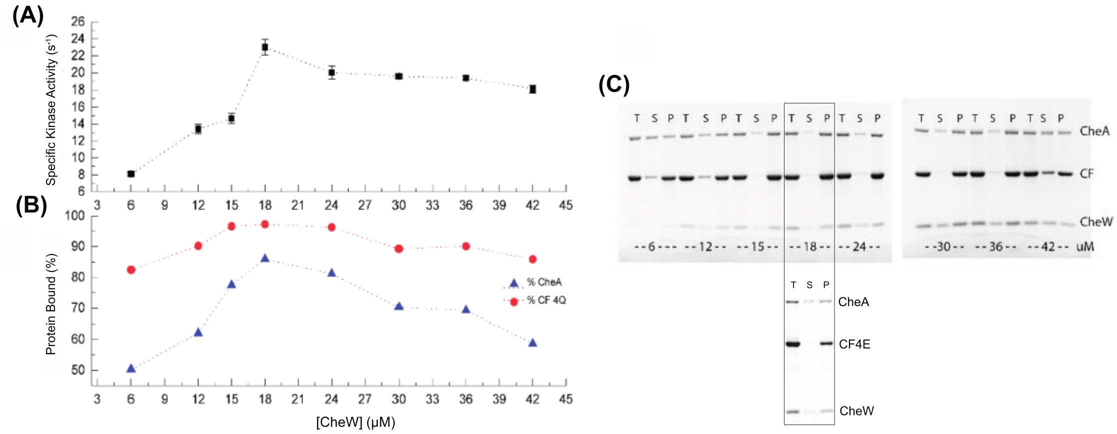


**Fig. S2.** Optimization of complex assembly conditions. 18 µM CheW exhibited the highest activity and highest percentage of CheA bound. (*A*) Specific kinase activity per total CheA of CF4Q complexes prepared with 30 µM CF4Q, 3 µM CheA, 860 µM lipid and varying CheW concentrations. Activity measurements are from three independent replicates. (*B*) Bound fraction of CF and CheA was determined by sedimentation assay. (*C*) SDS-PAGE of sedimentation assay: each lane is marked T (total), S (supernatant), or P (pellet). Percent bound was calculated as 100*(Total – Super)/Total. *Top*: gels for CF4Q complexes, corresponding to the optimization shown in *(B)*. *Bottom*: gel for CF4E under the chosen 18 µM CheW condition.

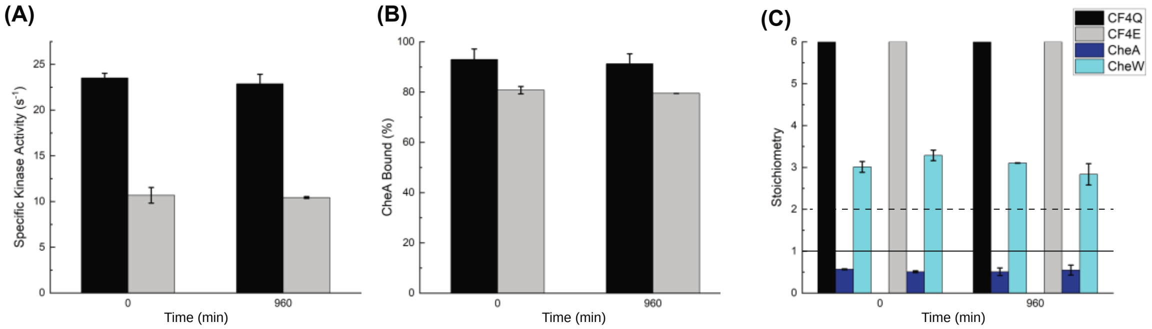


**Fig. S3.** CF complexes retain kinase activity and stoichiometry over the time-course of deuterium labeling. (*A*) Specific kinase activity and (*B*) Percent CheA bound in CF4Q complexes (black) and CF4E complexes (gray) at 0 min and 960 min. CF4Q and CF4E complexes activate CheA by about 600-fold and 250-fold, respectively (activity of CheA in solution ~ 0.04 $s^{-1}$). (*C*) Stoichiometry determined from sedimentation assays of CF4Q complexes and CF4E complexes at 0 min and 960 min. Native arrays have CF:CheA:CheW = 6:1:2. Chosen conditions with limiting CheA (blue) and excess CheW (Cyan) lead to a CF:CheA:CheW = 6:0.5:3. CheA level (solid line) and CheW level (dashed line) observed in native arrays.

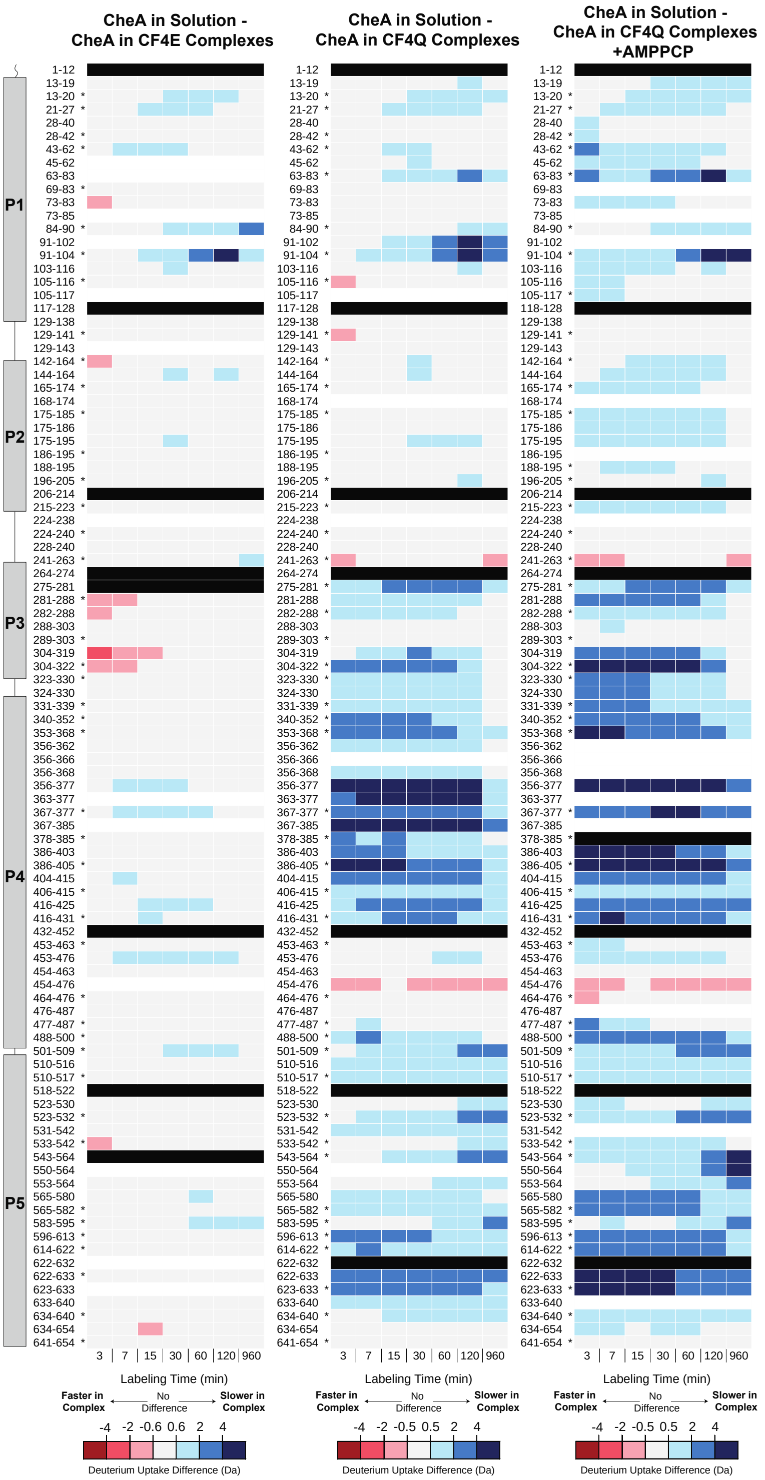


**Fig. S4.** Uptake difference between CheA in Solution and CheA in functional signaling complexes for all peptides and time points. Uptake differences for CF4E complexes (*left*), CF4Q complexes *(middle)* and CF4Q complexes with 10mM AMPPCP (*right*). Asterisks indicates peptides chosen for representing uptake and uptake differences on CheA structures in Figures 2-4. Black bars indicate regions with no coverage; white gaps indicate missing peptide, but other peptides provide coverage.

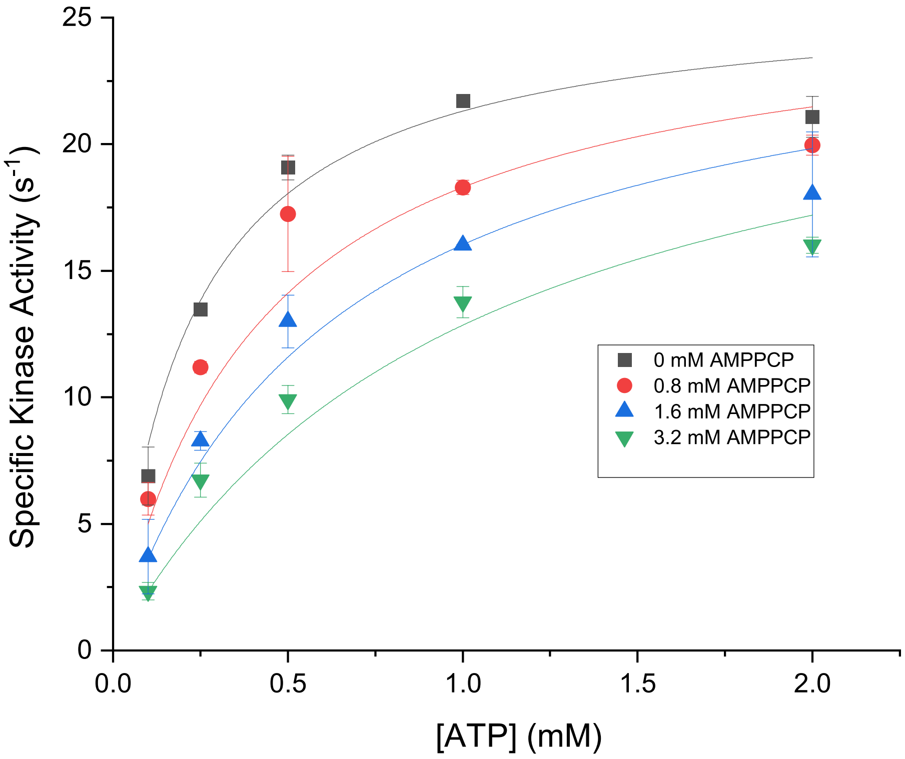


**Fig. S5.** CheA inhibition by AMP-PCP binding. Kinase activity was performed using limiting CheA conditions in the kinase-on state ((3 µM CheA, 18 µM CheW, 30 µM CF4Q, and 860 µM lipid) ) at 0.1, 0.25, 0.5, or 1.2 mM of ATP with varying amounts of AMP-PCP present (n=2). Competitive inhibition fitting was performed using Origin 2020 to determine Ki =877 µM for AMP-PCP.

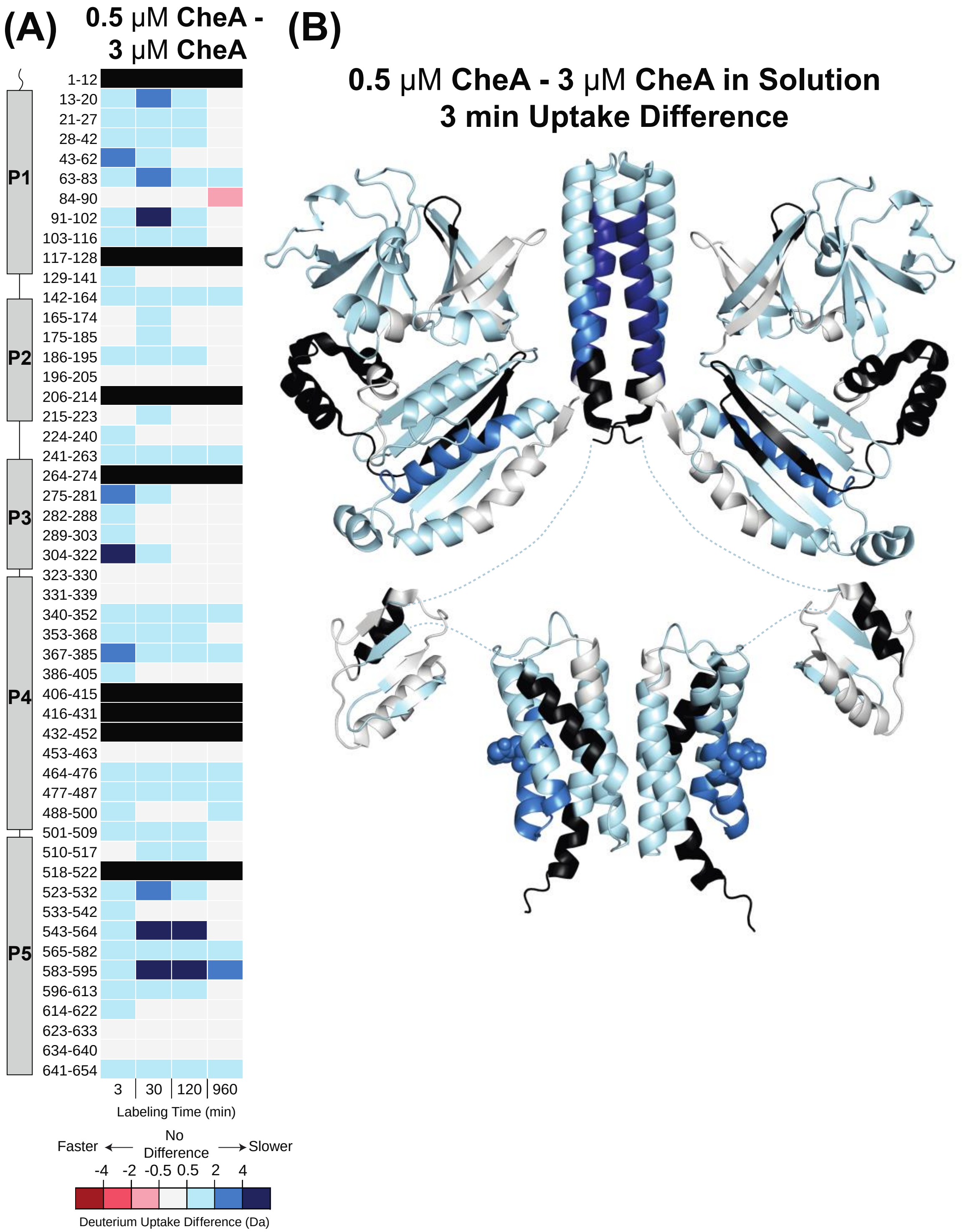


**Fig. S6.** Deuterium uptake differences due to increased dimerization of CheA in Solution. (*A*) Deuterium difference was calculated as described above. Red indicates faster exchange, gray indicates no significant difference, and blue indicates slower exchange of 3 uM CheA (75% dimer) relative to 0.5 uM CheA (50% dimer). Black indicates regions of no coverage. (*B*) Difference in uptake at 3 min is mapped onto CheA dimer model.

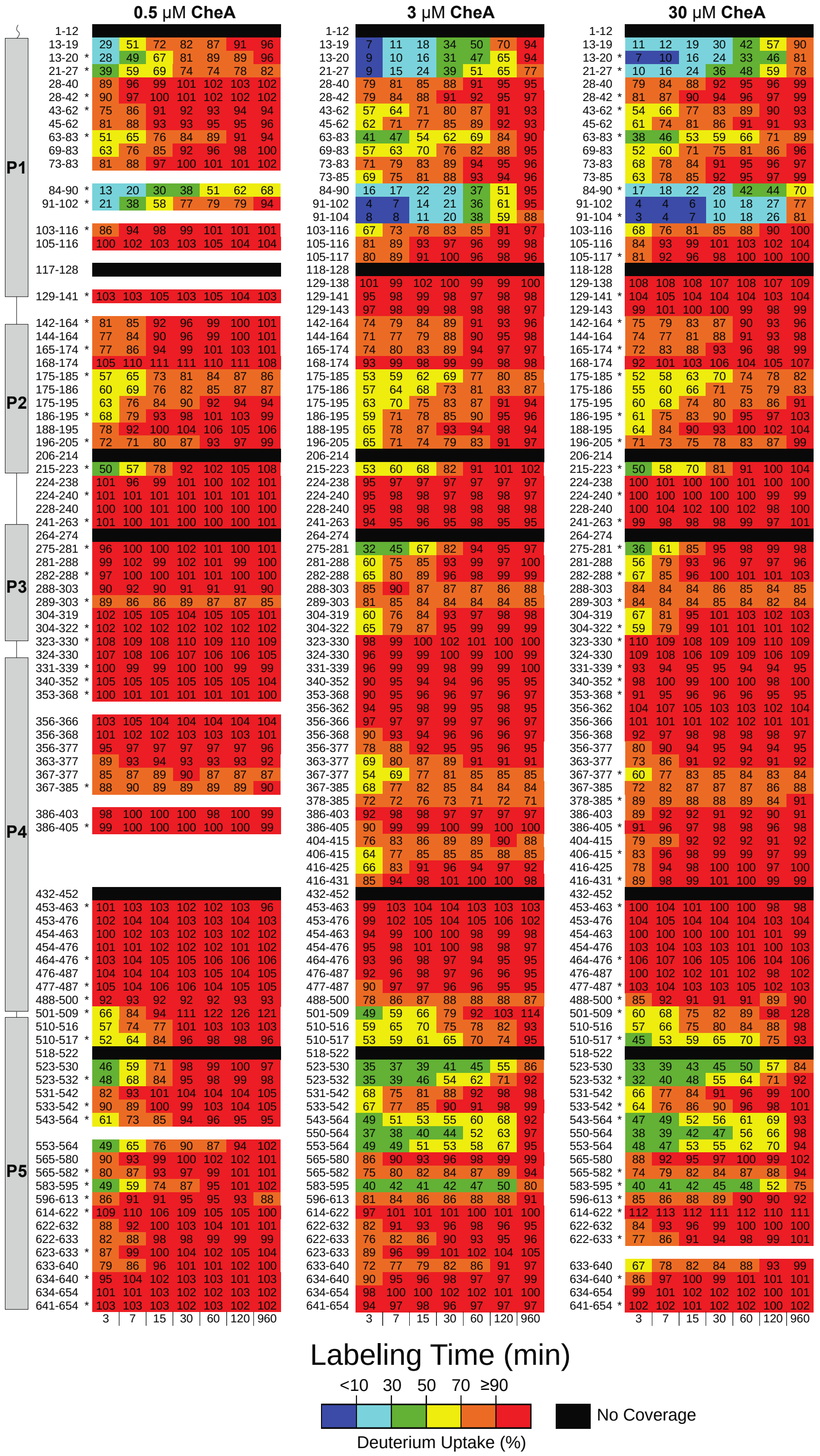


**Fig. S7.** Percent deuterium uptake of CheA in solution depends on fraction of dimer present. Uptake percentage of (*left*) 0.5 μM CheA (50% dimeric), (*middle*) 3 μM CheA (75% dimeric) and (*right*) 30 μM CheA (91% dimeric). Asterisks indicates peptides chosen for representing uptake differences on CheA structures in Figures 5 and S6. Black bars indicate regions with no coverage; white gaps indicate missing peptide, but other peptides provide coverage.

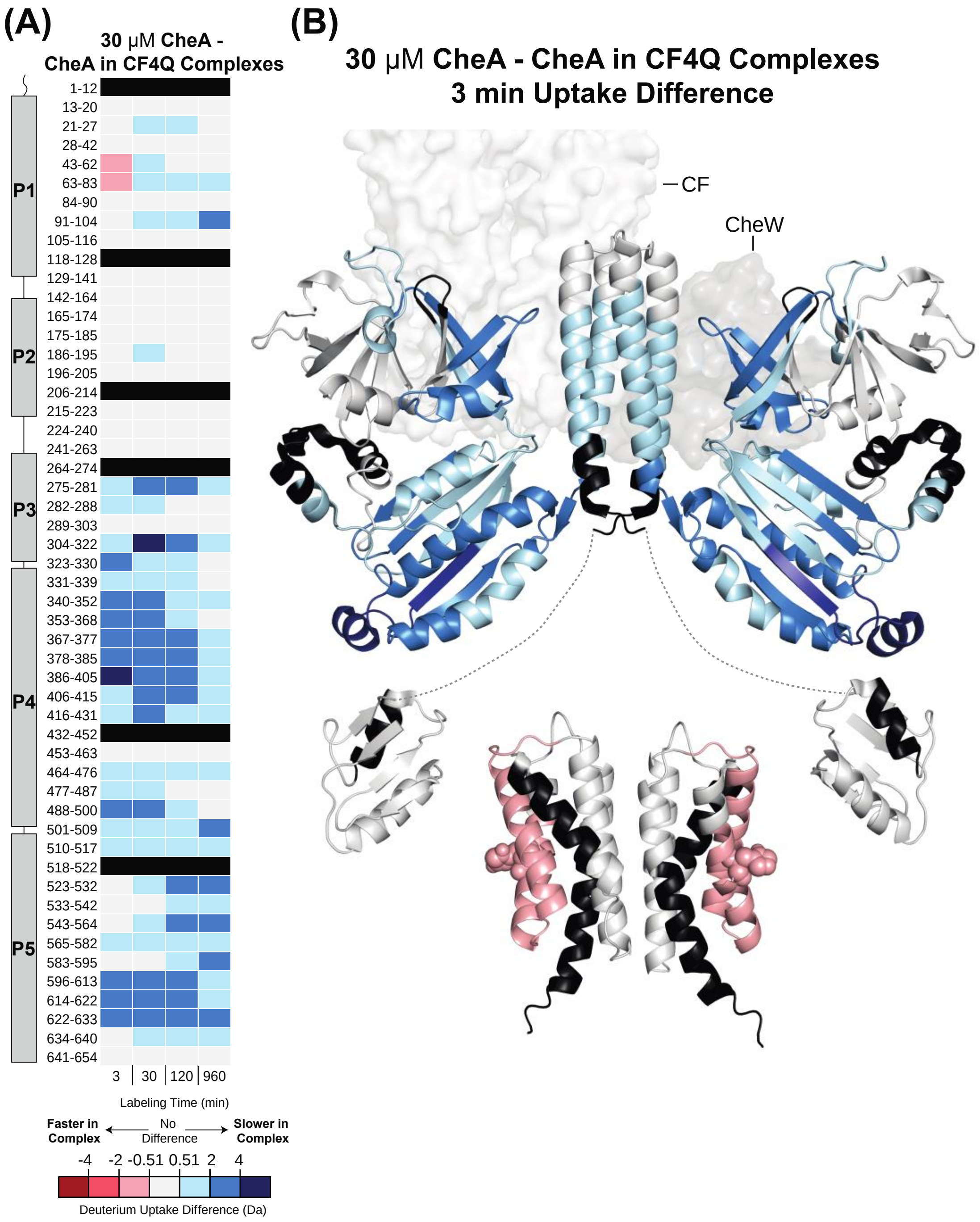


**Figure S8.** Deuterium uptake difference between dimerized CheA in solution and CheA in kinase-on signaling complexes. (*A*) Uptake difference for each peptide: uptake of 30 μM CheA (~91% dimer) minus uptake of CheA in CF4Q complexes. Difference uptake scale ranges from red indicating faster exchange of CheA in complexes to gray indicating no significant difference to blue indicating slower exchange in complexes. Regions with no identified peptide are indicated in black. (*B*) 3 min uptake difference is represented on the structural model of CheA as in previous figures.

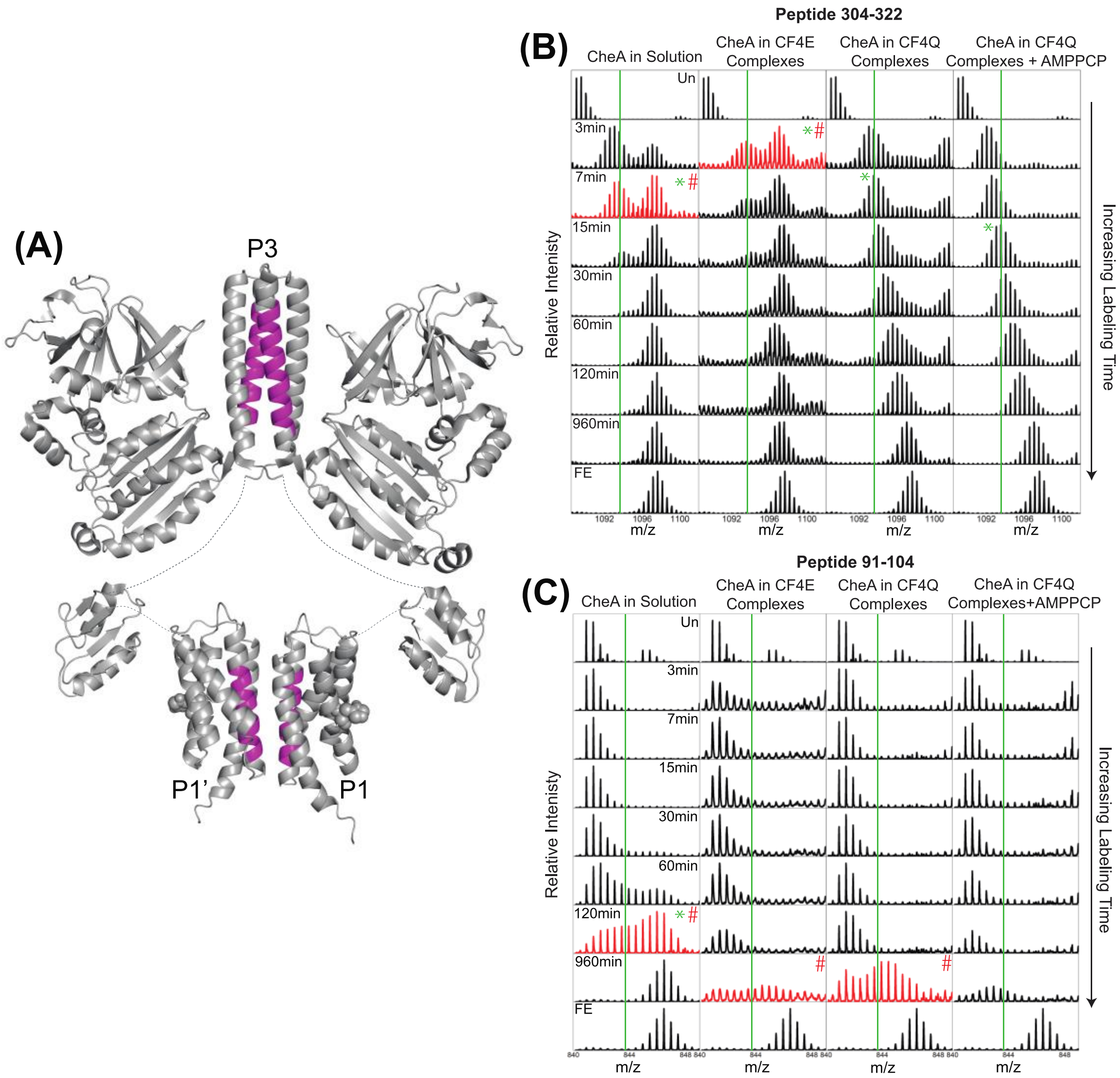


**Fig. S9.** Estimated $t_{1/2}$ of EX1 and EX2 at P3 and P1 dimerization interfaces. (*A*) Structural model of CheA dimer, highlighted in magenta to show peptides that exhibit EX1 exchange within the P3 and P1 dimerization domains. Stacked spectra (for one charge state in one replicate) of (*B*) P3 peptide 304-322, and (*C*) P1 peptide 91-104, in free CheA and in signaling complexes for unexchanged (Un), exchange timepoints, and fully exchanged (FE) samples. The $t_{1/2}$ for the EX1 process (indicated by red spectra marked with #) is estimated as the time at which the fully exchanged isotopic distribution has at least 50% of the total intensity. The $t_{1/2}$ for the EX2 process (indicated by a green asterisk) is estimated as the time at which the slower exchanging isotopic distribution has reached half of full exchange (vertical green line).

### (A) P1 peptide 91-102

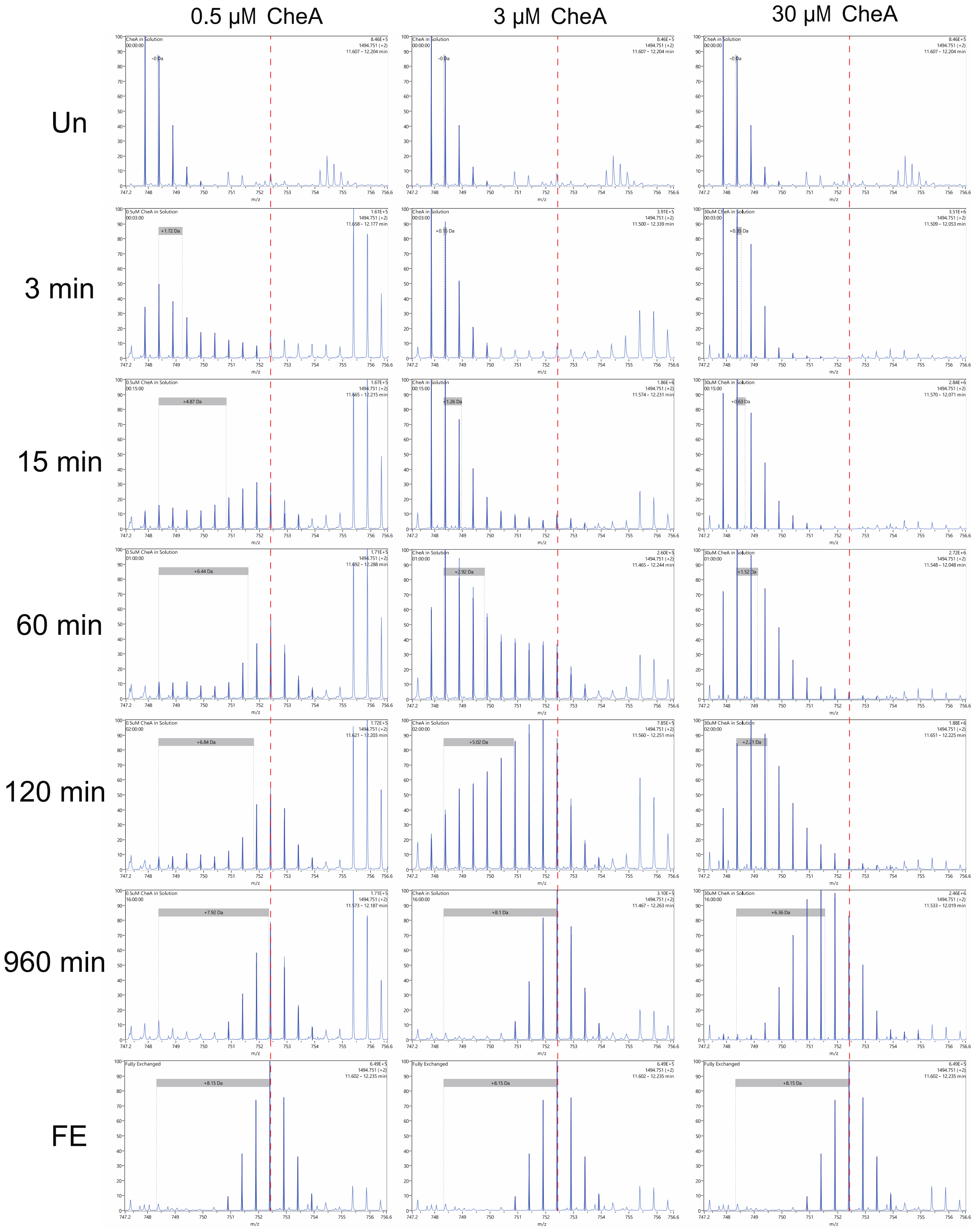

### (B) P3 peptide 304-322

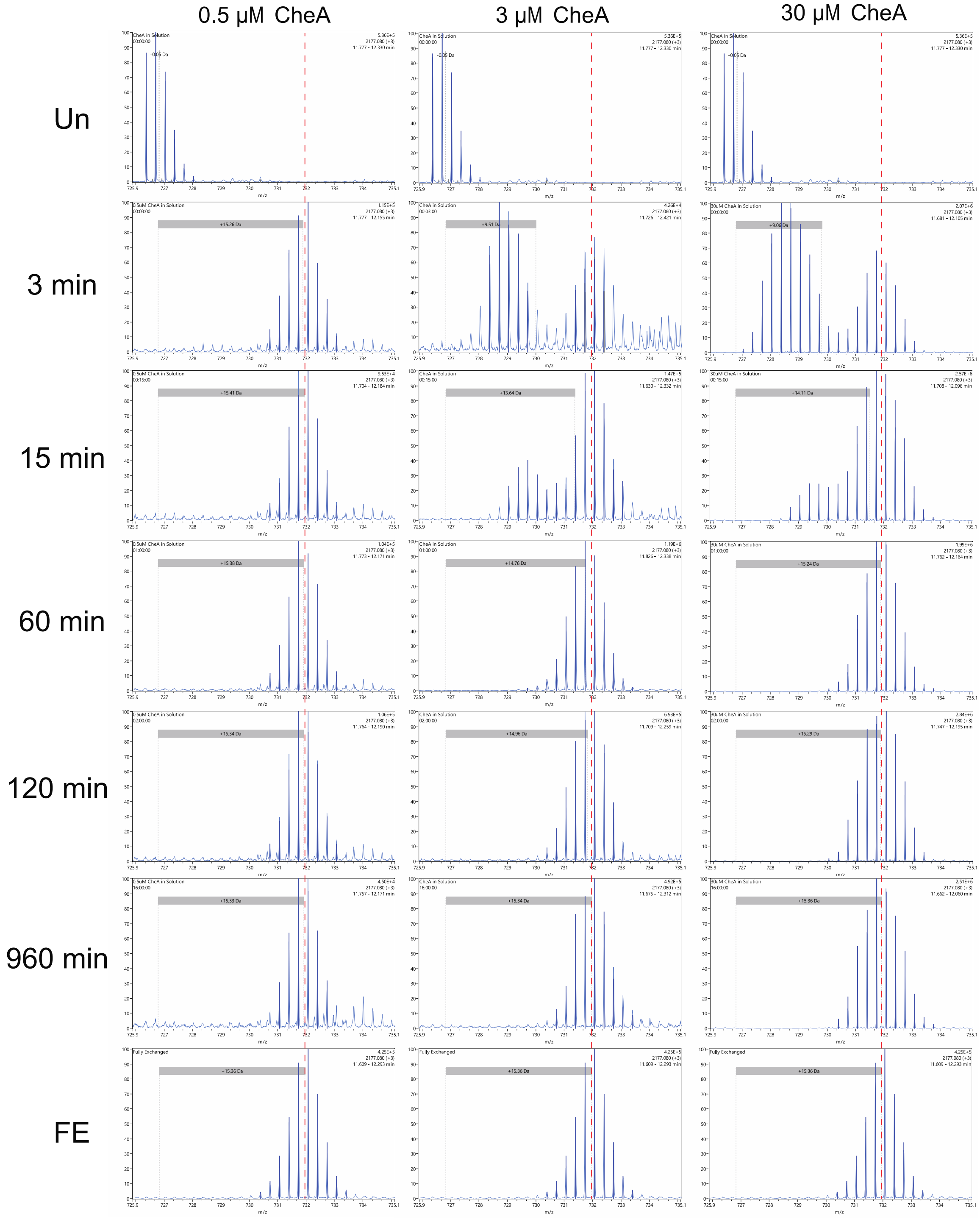


**Fig. S10**. Bimodal HDX patterns for dimer interface peptides of CheA vs % dimer. *(A)* P1 peptide 91-102. *(B)* P3 peptide 304-322. Spectra shown for one charge state in one replicate for each sample; red dashed line represents the center mass of the fully exchanged CheA sample for the displayed peptide. 0.5 μM CheA is 50% dimer; 3 μM CheA is 75% dimer, and 30 μM cheA is 91% dimer.

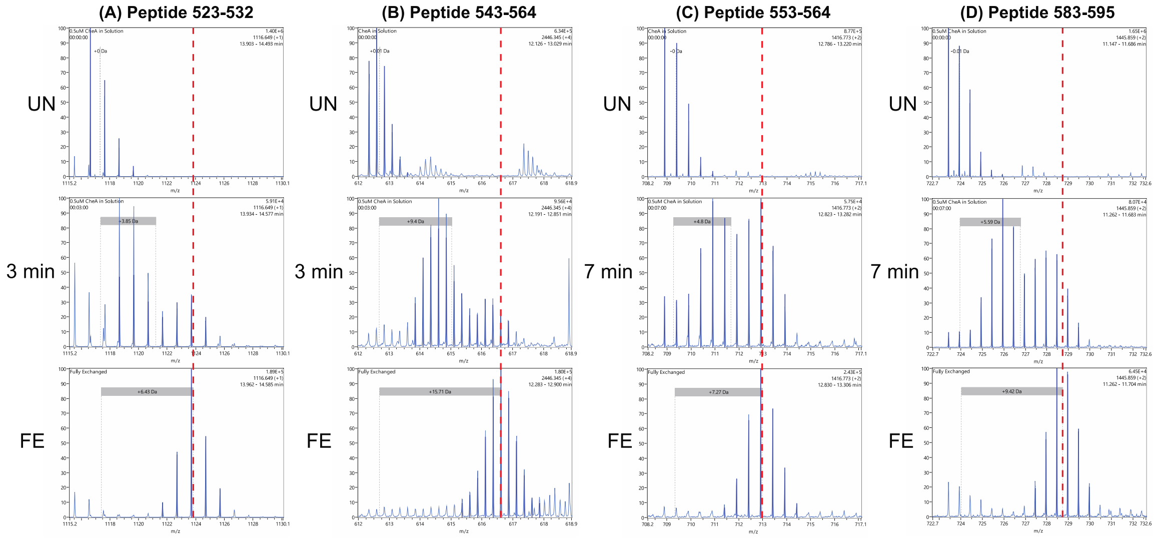


**Fig. S11**. Representative mass spectra illustrating bimodal HDX patterns observed in P5 peptides in 0.5 µM CheA (50% monomer) sample. Spectra shown for one charge state in one replicate for each peptide; red dashed line represents the center mass of the fully exchanged CheA sample for the displayed peptide.

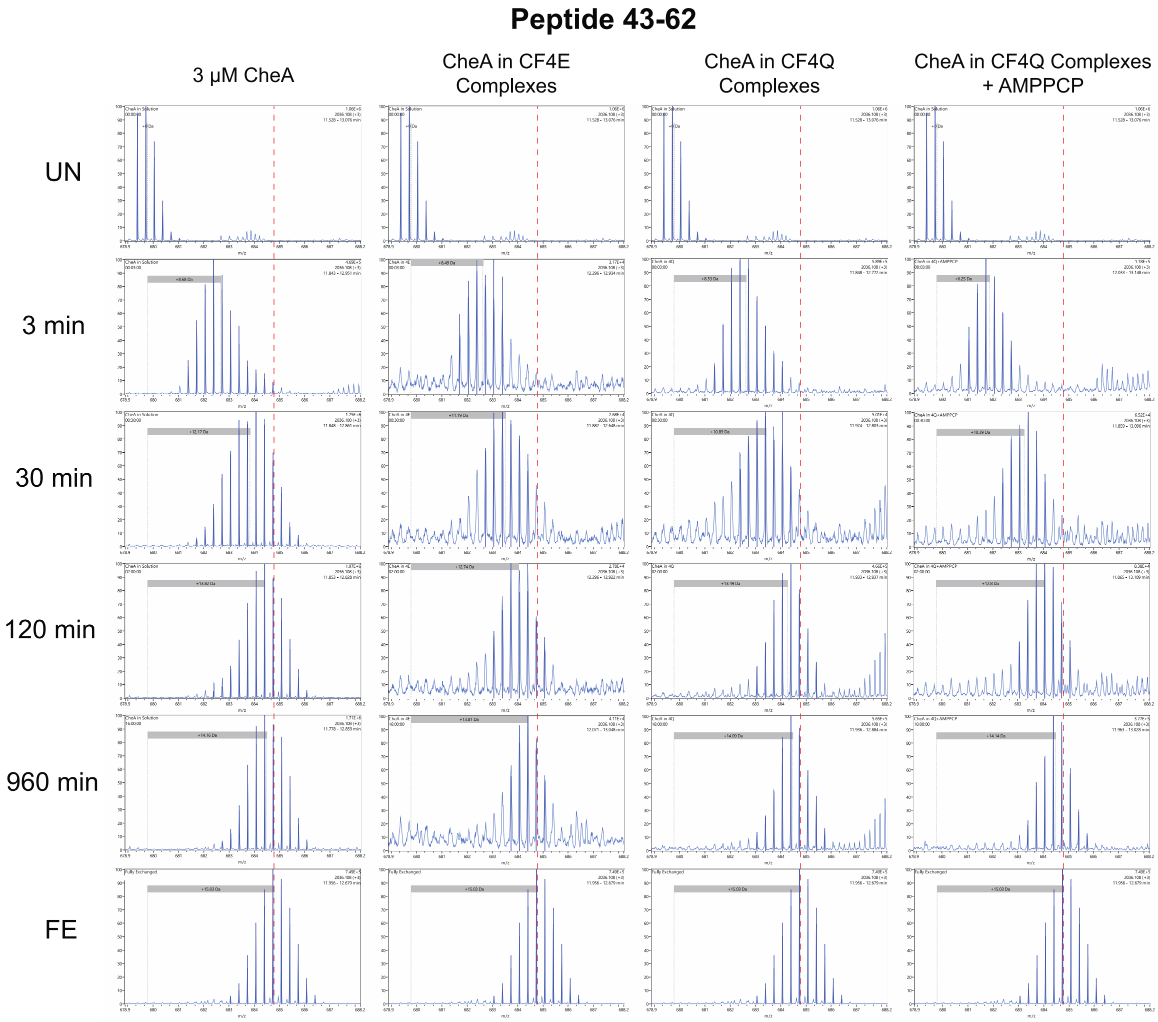


**Fig. S12**. Mass spectra illustrating HDX of peptide containing the H48 phosphorylation site. Spectra shown for one charge state in one replicate for each sample; red dashed line represents the center mass of the fully exchanged CheA sample for the displayed peptide.

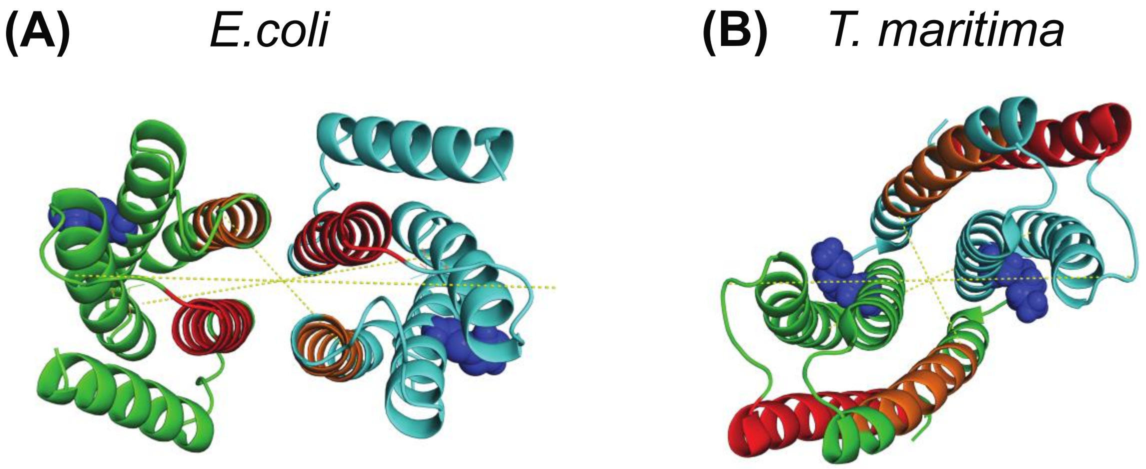


| | ***E. coli* model** | ***T. maritima* 1tqg** |
|---|---|---|
| Helix A (orange) | 13-27 | 10-24 |
| Helix D (red) | 84-104 | 81-101 |
| pds distance 25 Å | E15 (23.3 Å*+7=30.3) | E12 (16.3 Å*+7=23.3) |
| pds distance 30 Å | T66 (29.7 Å*+7=36.7) | C63 (24.4 Å*+7=31.4) |
| pds distance 62 Å | E80 (42.6 Å*+7=49.7) | E77 (38.7 Å*+7=45.7) |
| Phosphorylation site (blue) | H48 | H45 |

*Cα to Cα

**Fig. S13.** Comparison of P1 dimer models. *(A)* Proposed model for *E. coli* P1 dimer based on slow HDX of helices A and D (this study). *(B)* Crystal structure (1tqg) of *T. maritima* P1 dimer (9). Residues with slow HDX in signaling complexes of *E. coli* CheA are colored orange (helix A) or red (helix D). Dashed lines indicate distances measured by pulsed dipolar EPR spectroscopy (pds) on signaling complexes containing *T maritima* CheA (9); pds distances are in reasonable agreement with both models.

**Table S1.** Changes in covalent labeling (CL) and HDX in P3P4P5 upon incorporation of CheA into kinase-on complexes with CF4Q.

| Domain | Residue | ΔCL[1] | ΔHDX[2] | Interpretation |
|---|---|---|---|---|
| P4 | Y331 | decrease | decrease | Δ solvent accessibility |
| P4 | T375/H376 | none | decrease | Δ dynamics |
| P4 | S381/H384 | decrease | decrease | Δ solvent accessibility |
| P4 | S399 | none | decrease | Δ dynamics |
| P4 | H488 | none | decrease | Δ dynamics |
| P5 | H543 | decrease | decrease | Δ solvent accessibility |
| P5 | Y614 | none | decrease | Δ dynamics |
| P5 | K616 | none | decrease | Δ dynamics |

[1]Complex vs 51 µM CheA in solution (93% dimer). Data from reference (10).
[2]Complex vs 3 µM CheA in solution (75% dimer).
